# Uncovering Bifurcation Behaviors of Biochemical Reaction Systems from Network Topology

**DOI:** 10.1101/2024.11.19.624026

**Authors:** Yong-Jin Huang, Takashi Okada, Atsushi Mochizuki

## Abstract

The regulation of biological functions is achieved through the modulation of biochemical reaction network dynamics. The diversity of cell states and the transitions between them have been interpreted as bifurcations in these dynamics. However, due to the complexity of networks and limited knowledge of reaction kinetics, bifurcation behaviors in biological systems remain largely underexplored. To address this, we developed a mathematical method, Structural Bifurcation Analysis (SBA), which decomposes the system into substructures and determines important aspects of bifurcation behaviors—such as substructures responsible for bifurcation conditions, bifurcation-inducing parameters, and bifurcating variables—solely from network topology. We establish a direct relationship between SBA and classical bifurcation analysis, enabling the study of systems even in the presence of conserved quantities. Additionally, we provide a step-by-step bifurcation analysis for general use. We applied our method to the macrophage M1/M2 polarization system. Our analysis reveals that the network structure strongly constrains possible patterns of polarization. We also clarify the dependency of the M1/M2 balance on gene expression levels and predict the emergence of intermediate polarization patterns under gene deletions, including SOCS3, which are experimentally testable.

## 1 Introduction

Various biological systems have been understood by employing mathematical approaches. Dynamical behaviors of biological systems are understood from interactions between biomolecules using mathematical models. For example, diversity of cell types and transition between them have been understood as bifurcation of dynamics of biological systems. Bifurcation is a phenomenon where the equilibria (or, more generally, the attractors) of a dynamical system undergo qualitative changes in response to continuous changes in system parameters.

Accumulated evidence has shown the connections between bifurcation theory and biological phenomena. For instance, Ozbudak et al. found that in *Escherichia coli*, the expression level of lactose permease (LacY), which is responsible for the cellular uptake of lactose and its analogues (e.g. TMG), switched significantly in response to continuous changes in the nutrient environment [1]. Ozbudak et al. developed a simple mathematical model and found that the jumps in LacY expression can arise from transitions between different steady states associated with the passage of a saddle-node bifurcation point. In this case, not only does the bifurcation create distinct cellular states, but it also creates a situation in which the transitions between these states are reversible, but the nutrient concentration thresholds that trigger the transitions are different in a state-dependent manner.

Another biological example associated with bifurcation is the polarization of macrophages. Macrophage is a type of white cell that can be polarized toward either an M1 (pro-inflammatory) or an M2 (anti-inflammatory) phenotype [2]. M1 and M2 macrophages can coexist under some microenvironmental conditions, which indicates that the dynamics of the polarization features the existence of alternative steady states [3]. Moreover, it has been known that a macrophage has the ability of switching its phenotype in response to stimuli [4, 5, 6, 7]. These facts indicate that the dynamics of macrophage polarization allow transition between multiple steady states in response to parameter changes.

Bifurcation analyses of biological systems are mostly done in a heuristic manner. The analysis first starts from the construction of a mathematical model for the system under consideration, and then it is followed by the identification of equilibria in correspondence to given parameters. For a biological system, an analytical solution to determine the equilibrium point(s) by a function of parameters is not always technically available. Hence, to probe the bifurcation of a complex system, the equillibrium points are numerically determined while a parameter of the system is varying. This profiling step may capture bifurcation phenomena by chance; nonetheless, it is usually not known in advance that change in which of the numerous parameters can trigger a bifurcation. Also, it is not known in advance under what parameter condition bifurcation will occur, or which of the many variables will exhibit bifurcation behaviors.

The Jacobian matrix, denoted by *J*, plays a crucial role in the study of equilibrium bifurcations, as it takes the value of zero at a bifurcation point. Exploiting this property, a series of studies has provided fruitful findings that correlate the graphic representation of the network and the Jacobian determinant [8, 9, 10]. In addition, it has been shown that the number of alternative states can be even concluded from the sign of the Jacobian determinant [11, 12]. While the Jacobian matrix can help identify the onset of a bifurcation, a numerical approach is still required unless the matrix is explicitly defined as a function of parameters. In practice, for large networked systems, conducting a bifurcation analysis remains challenging.

To cope with the practical problems of bifurcation analysis rooted in the application of Jacobian determinant, Okada et al. developed Structural Bifurcation Analysis (SBA) by which the equilibrium bifurcation of a chemical reaction system is analyzed only from the structural information of the network [13, 14]. Structural Bifurcation Analysis is based on a matrix ***A*** determined by the network structure and a proved fact that det ***A*** = 0 only when det *J* = 0. This approach circumvents the problem that, without the functional form of reaction rate functions, the equilibria are not explicitly available for a bifurcation analysis. Nonetheless, a great deal about the possible bifurcation behaviors of the steady state can actually be determined from the network structure alone. Specifically, (1) By SBA, a network is decomposed into subnetworks based on a characteristics/index of sub-graphs, and this decomposition indicates all the conditions under which bifurcation can occur. (2) For each substructure given by the network decomposition, we can identify reactions containing the parameters that can induce the bifurcation. (3) Moreover, we can identify chemicals on the network that exhibit bifurcation behaviors when any substructure satisfies the bifurcation condition (Fig. 1).

**Figure 1:**
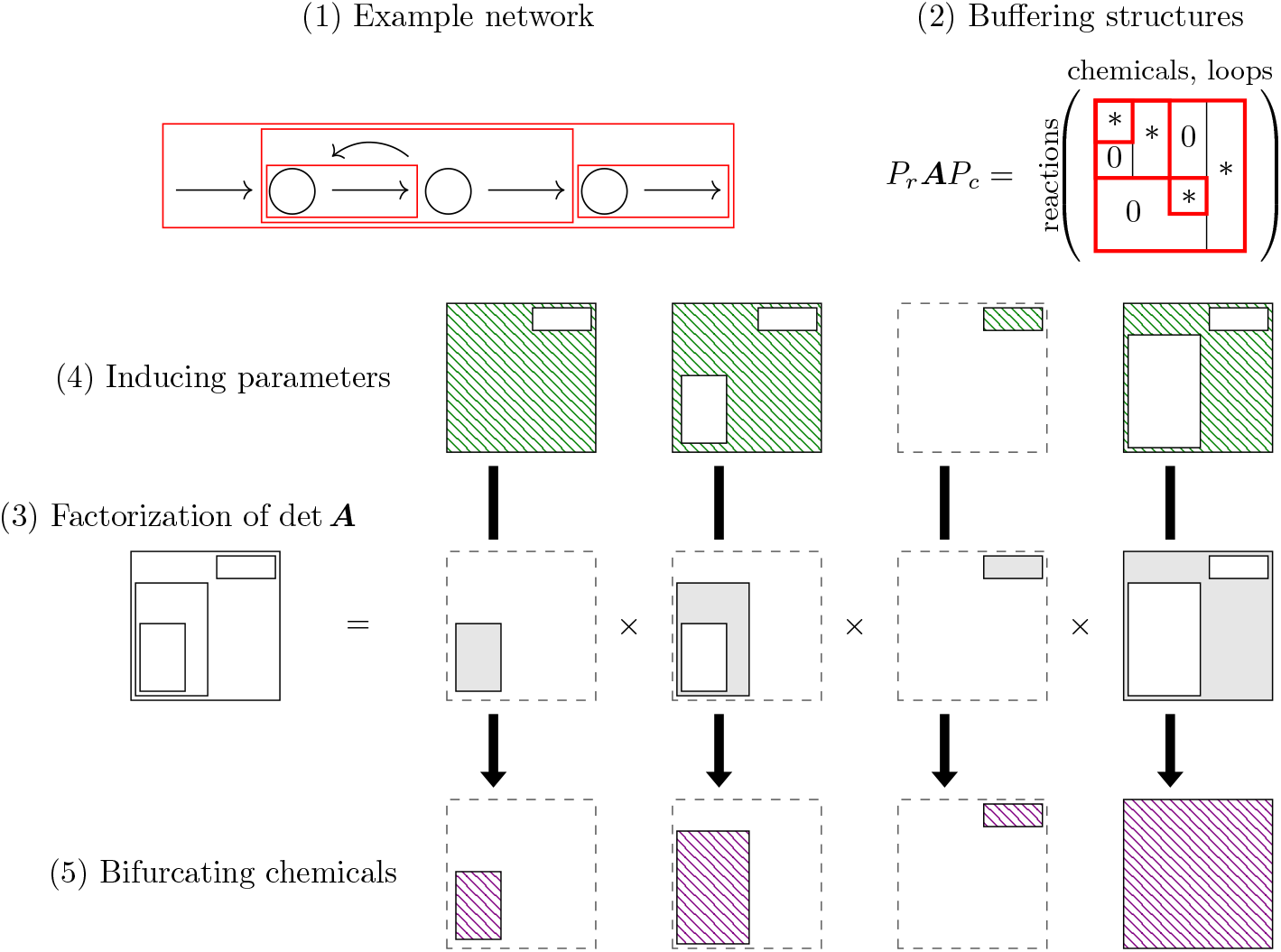
Summary of Structural Bifurcation Analysis (modified from [13]). One can construct a matrix ***A*** according to the network structure of a given chemical reaction networks. With the matrix, one can further conclude the conditions where bifurcations can occur, the inducing parameters, and the chemicals that exhibit the bifurcations.

That being said, there remained a couple of open problems in this approach: (i) the theorem was proved under the assumption that no conserved quantity is present in a given network, whereas the general case is open to discussion; (ii) the explicit relation between the matrix ***A*** and Jacobian matrix is unclear, giving rise to a barrier to further development and applications. In this paper, we extend the theory of SBA by solving the two open problems. Our new theorem provides an explicit formula converting ***A*** to a modified Jacobian matrix *J*_*g*_ (without redundant eigenvalues arise from the presence of conserved quantities, see Method). This breakthrough not only guarantees the application of SBA in general cases, but it also gives a corollary as a criterion for the (in-)stability of equilibrium points. Remarkably, our theorem is kinetics-independent and renders study of system dynamics a hierarchical insight into network topology; furthermore, it provides a great potential to associate the framework of structural analysis with other theorems rooted in Jacobian matrices, e.g., multistationarity analysis.

## 2 Results

### 2.1 Unveiling bifurcation behaviors of chemical reaction networks through SBA

We begin by demonstrating how bifurcation behavior in chemical reaction networks can be predicted using SBA based on network topology. Suppose that a network composed of four reactions is given:

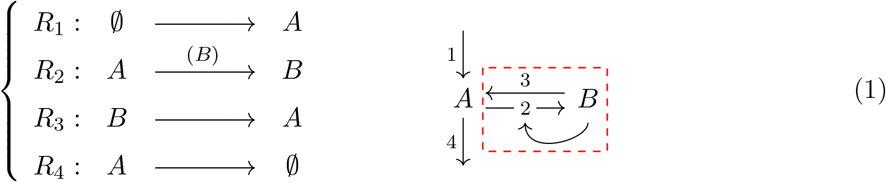

Here, the symbol ∅ denotes an external reservoir. The arrow “→” from chemical *B* to reaction 2 in the network graph represents positive regulation. The subnetwork indicated by the red dashed block is a *buffering structure* identified by SBA, of which definition and properties are explained later. Then, according to the differences in the coefficients in the chemical reactions, we know the system follows the ordinary differential equation

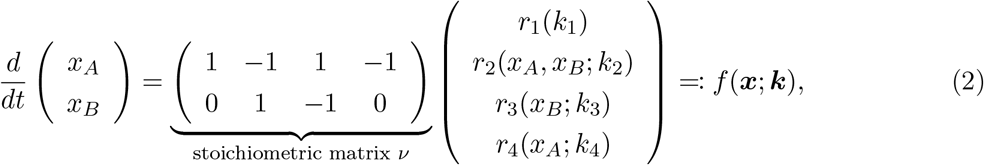

where *k*_*n*_’s are parameters of the reaction rate functions.

The SBA uses only the structural information from a reaction system, which is encapsulated in an augmented matrix ***A*** (see Method Eq. (15) for the precise definition). In the absence of conserved quantities, ***A*** is constructed based on the information whether reaction rates depend on each chemical (i.e., 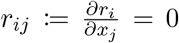 or not) and a basis for the right kernel of the stoichiometric matrix ν. A kernel basis can be found from a perspective of graph theory by considering *closed paths/loops*, with the reservoir ∅ also taken as a node of the graph. In this case, we observe two loops: 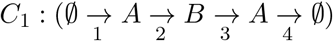 and 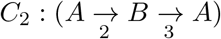. Then, it is immediately followed by

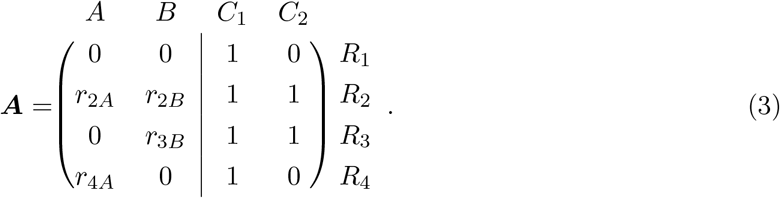

By the theorem given by Okada, Tsai, and Mochizuki [13], an equilibrium bifurcation occurs only when the determinant of the matrix takes the value of zero, since det ***A*** ∝ det *J*_*f*_ in which *J*_*f*_ is the Jacobian matrix of *f* in the governing equation Eq. (2). Therefore, the bifurcation points that can be identified by the condition det *J*_*f*_ = 0 can also be identified by det ***A*** = 0.

One of the advantages of SBA over standard analysis using Jacobian matrix is that it gives a network decomposition with the factorization of bifurcation conditions. A network can be decomposed when there exist buffering structures [15, 16], defined to be a subnetwork *γ* satisfying:

1. (Output-completeness) All the reactions regulated by any chemicals of *γ* are included in *γ*.
2. 0 = #(chemicals in *γ*) − #(reactions in *γ*) + #(loops in *γ*) + #(conserved quantities in *γ*).

This is followed by a fact (which can be an alternative definition, see Appendix A: Buffering structures and the localization principle) that the matrix ***A*** can be rearranged to be an upper block triangular matrix; that is, for some permutation matrices *P*_*r*_ and *P*_*c*_ we have

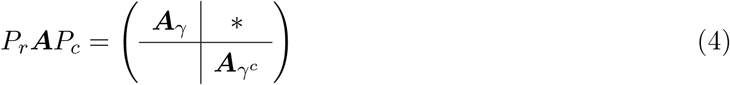

such that ***A***_*γ*_ is a square block, with *γ*^*c*^ denoting the complement of *γ*. The immediate consequence of the above factorization property is that a bifurcation occurs only if either (i) det ***A***_*γ*_ = 0 or (ii) det 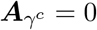 For the considered example, *γ* := {*B, R*_2_, *R*_3_} is a buffering structure and one has

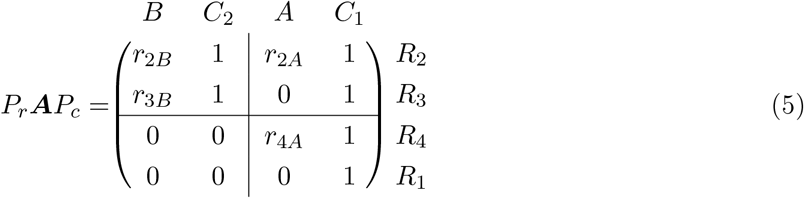

with the factorization of bifurcation conditions

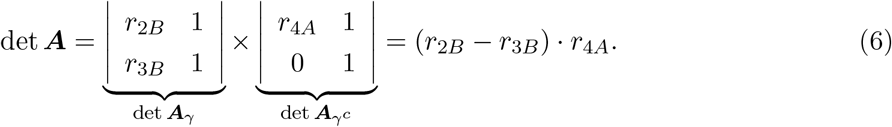

and it is clear that det 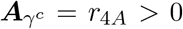 because *A* is a reactant of *R*_4_; therefore, we can conclude that the system bifurcates only when det ***A***_*γ*_ = *r*_2*B*_ − *r*_3*B*_ = 0.

Another advantage of SBA is its ability to predict bifurcation behavior without requiring kinetic information. With SBA [14], it can be concluded that (i) when det ***A***_*γ*_ = 0, only chemicals in *γ* would exhibit the bifurcation behavior (Appendix A: Buffering structures and the localization principle); (ii) if det 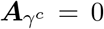, all the chemicals could exhibit the bifurcation behavior. Nonetheless, in the considered example, we see that a bifurcation occurs only when det ***A***_*γ*_ = *r*_2*B*_ − *r*_3*B*_ = 0, and hence the behavior would only be exhibited by the chemical *B*. To verify that theoretical prediction holds regardless of kinetics, we plot the bifurcation diagrams of the common example network with three different reaction functions in ordinary differential equation models (Appendix B: the models used for method demonstration). With a parameter of the rate function of reaction *R*_2_ is changing, the network exhibits three different types of bifurcations, whereas the bifurcation behaviors are always exhibited only by *B* as predicted (Fig. 2).

**Figure 2:**
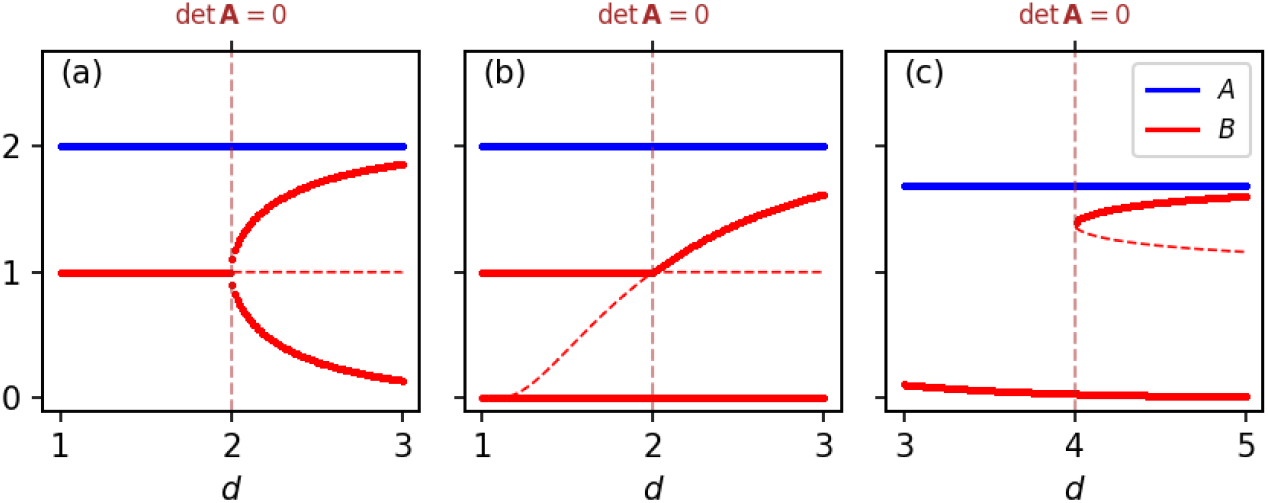
Bifurcation diagrams of the example network with different ODE models. The three models respectively show a (a) pitchfork, (b) transcritical, and (c) saddle-node bifurcation. Independent of the kinetics, we verify our prediction that the bifurcation behavior is exhibited only by *B*. The ODE models and the parameter settings are given by Appendix B: the models used for method demonstration.

### 2.2 The generalization of SBA with a standard procedure for chemical reaction networks

One of the challenges in bifurcation analyses in analytical approaches is posed when a system possesses conserved quantities. Conserved quantities create problems for standard bifurcation analysis because they cause the determinant of the standard Jacobian to vanish in such cases (as discussed in Method). Zero eigenvalues associated with these conserved quantities make det *J*_*f*_ ≡ 0, rendering the determinant a poor indicator of bifurcation. Networks with conserved quantities have been studied in the structural analysis with an extended ***A*** with data on conserved quantities, but not in the context of bifurcation analyses but sensitivity analyses [16]. It was left as an open question whether such a structural approach still works for exploring bifurcation phenomena, that is, whether det ***A*** acts as a non-trivial bifurcation indicator despite the failure of det *J*_*f*_.

Facing this problem, we find that the theory of SBA can be generalized to the case in which conserved quantities are present. Specifically, we prove that when conserved quantities are present, det ***A*** is proportional to a *modified* Jacobian *J*_*g*_ that omits the redundant eigenvalues linked to conserved quantities. In other words, det ***A*** remains a reliable indicator of equilibrium bifurcations even in systems with conserved quantities (Method Eq. (18) and Eq. (19)). In particular, an explicit formula that transforms the matrix ***A*** into the modified Jacobian *J*_*g*_ is given by our main theorem (**Theorem 1** in Method), followed by an identity

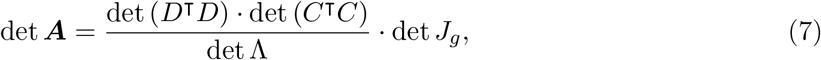

where the constant matrices *C, D*, and Λ are structurally determined and defined in terms of the stoichiometric matrix. Furthermore, the system stability is assessed by det ***A*** with

#### Corollary 1

sign(det Λ · det ***A***|_***x***_) ≠(−1)^*M*−*L*^ ⇒ ***x*** *is not a stable equilibrium point*,

where *M* is the number of chemical species, and *L* is the number of conserved quantities (see Method).

In summary, our new mathematical findings demonstrate that the determinant of the matrix ***A***

serves as a general bifurcation and stability indicator for chemical reaction networks. This extends the applicability of SBA to a broader range of networks, including the problematic cases in which the standard Jacobian *J*_*f*_ fails to be an informative indicator. Moreover, although the definition of ***A*** differs in the presence of conserved quantities, the procedure suggested by ref. [13] for cases without conserved quantities can still be formally adopted without modification. To further streamline the workflow of SBA, we introduce a 6-step procedure (Fig. 3):

**Figure 3:**
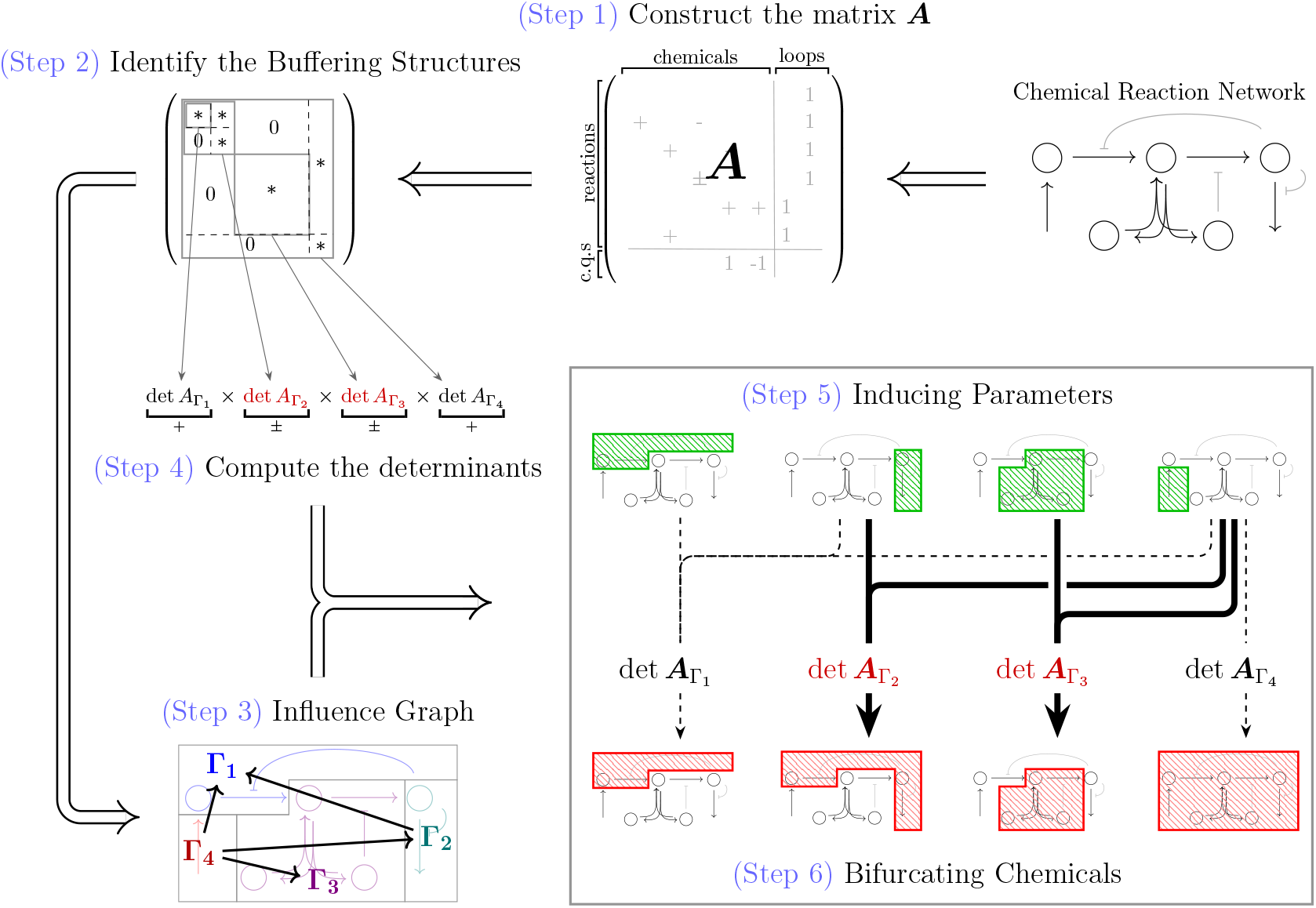
Summary of the standard procedure of Structural Bifurcation Analysis. The theory of SBA has been extended to networks with conserved quantities allowed, and we further improve the procedure for users to facilitate the roles and relations of subnetworks in the bifurcation dynamics.

1. Construct the matrix ***A*** according to the network topology.
2. Identify buffering structures and rearrange ***A*** into an upper block triangular form.
3. Decompose the network into subnetworks Γ_*i*_’s and construct the influence graph according to the rearranged ***A***.
4. For each subnetwork Γ_*i*_, compute det 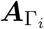
5. Summarize the conditions in which det 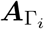 can change its sign for some Γ_*i*_, and conclude the inducing parameters.
6. For each condition listed in (5), according to the influence graph, determine the bifurcating chemicals.

We improve the protocol with the step of constructing the influence graph (defined in Method Eq. (28) and Eq. (29)), which provides an insight into the roles of and relations between subnetworks. With the generalization of the theory and the improvement of the procedure, in the following subsection, we would like to investigate the switching behaviors of a biological system at a cellular level.

### 2.3 Macrophage Polarization Network

Macrophages are specialized innate immune cells with significant plasticity to change their phenotypes in response to the microenvironment [17]. The phenotypic states of macrophages are broadly categorized into two distinct groups: M1 (pro-inflammatory) and M2 (anti-inflammatory), each wielding diverse functions in inflammation and tumor defense. M1 macrophages, characterized by highly activated STAT1 and NF*κ*B, produce pro-inflammatory cytokines and exert cytotoxic functions to combat pathogens [18, 19]; on the other hand, M2, characterized by highly activated STAT3 and STAT6, represents an anti-inflammatory group which takes charge of tissue repairing and contributes to the resolution of inflammation [18, 20, 21]. In various diseases, the balance between M1 and M2 macrophages can be crucial and is thought to be strictly regulated according to the microenvironment including physiological and pathological conditions. Thus, for therapeutic purposes, researchers strive to comprehend and potentially manipulate macrophage polarization. For instance, it has been suggested that mutations in SOCS3 (suppressor of cytokine signaling 3), a gene that has a suppressive function for STAT3, may result in a hyperactivation of STAT3 and thus prevent the metastasis of skin cancer [22]. By elucidating the macrophage switching system, it would be possible to artificially switch macrophages between different phenotypes, which could potentially be used in the future for immunotherapy against cancer.

We take a simplified network associated with macrophage polarization as an example (Fig. 4). With the network, we first numerically verify that ***A*** outperforms the standard Jacobian *J*_*f*_ as a bifurcation indicator (by **Theorem 1**) and a stability indicator (by **Corollary 1**). After that, we demonstrate the power of SBA on predicting bifurcation behaviors based on the network topology. It is noteworthy that a signaling pathway as considered can usually possess conserved quantities, because signal transduction pathways often contain signaling proteins that reversibly transition between activated and inactivated states, and the total amounts of such (paired) proteins are constant in the timescale of chemical reactions. Specifically, in this macrophage polarization system, the total amount of each protein molecule (STAT1, NF*κ*B, STAT3, and STAT6) in a single cell corresponds to a quantity that does not change in a short period of time. Note the difference between the total amount and the activity level of each molecule. An important fact is that the expression levels of these proteins are known to vary among cells. In particular, it is reported that STAT1-deficiency in macrophage leads to a significantly low tendency of switching to an M1 phenotype [23, 24]. Therefore, in the following analysis, we will focus on the qualitative changes in system behavior, or bifurcation behavior, that are caused by differences in expression levels between cells, i.e., differences in the total amount of signaling proteins (specifically, the total amount of inactivated and activated STAT1).

**Figure 4:**
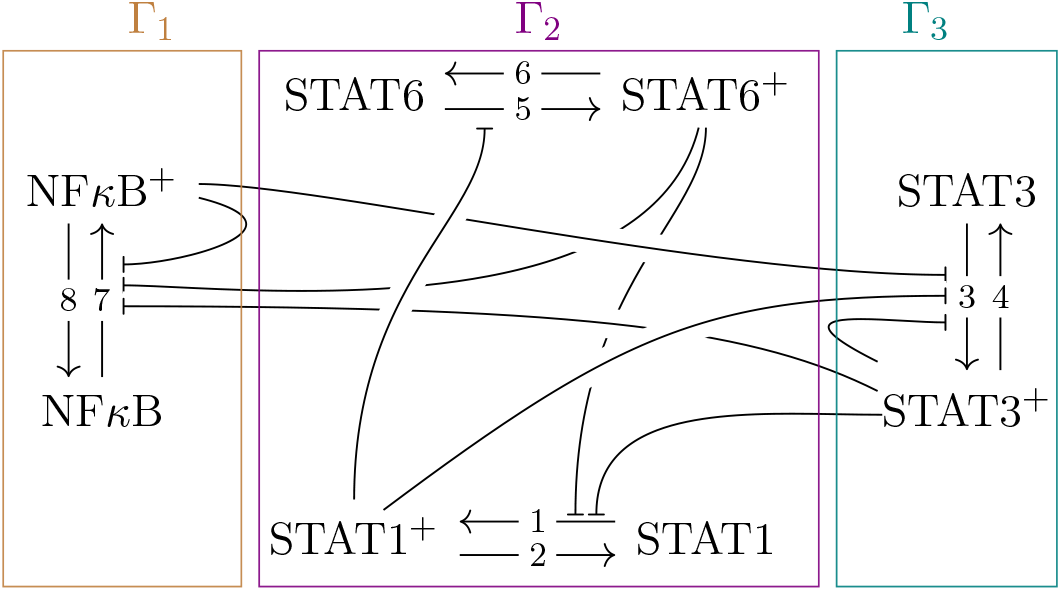
Macrophage polarization network for the wild type. By an arrow “⊣” pointing from a complex to a reaction, we mean a negative regulation. The superscript + denotes the phosphory-lated/activated form for each of the four complexes STAT1, STAT3, STAT6, and NF*κ*B, of which signalings are essential to the macrophage M1/M2 polarization. In this simplified network, the M1 polarization is primarily associated with the dominance of STAT1^+^ and NF*κ*B^+^, while the M2 polarization is associated with the dominance of STAT3^+^ and STAT6^+^.

#### 2.3.1 SBA for the (wild-type) macrophage polarization network

The network of the macrophage polarization system, presented in Fig. 4 and referred to as the wild-type (WT) network, possesses no buffering structures (except for a trivial one, the entire network). Consequently, our structural method implies that the whole network is responsible for the bifurcation condition, and the determinant factor associated with the entire network, det ***A***, is a bifurcation/stability indicator. Furthermore, it implies that if the system exhibits bifurcations, then all the four proteins must exhibit bifurcating behaviors.

To numerically verify these predictions from the SBA, we construct a model for the considered network. Reactions in the network can be categorized into two classes: deactivation (or degradation) and activation subject to negative regulation. For simplicity, the reaction rate function of deactivation is given by the law of mass action; on the other hand, the activation as well as the negative regulation involve phosphorylation, which is commonly modeled as a sigmoid function [25], and this is considered in our model as well. The detailed parameter settings for the following numerical experiments can be found in supplementary materials (Appendix B: the models used for method demonstration).

For the convenience of the notation, the concentration of NF*κ*B is denoted by *N*, and that of activated form NF*κ*B^+^ is denoted by *N*^*p*^. Similarly, *S*_*i*_ and 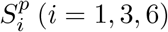 denote the concentrations of inactivated and activated STAT family, respectively. Then, the four conserved quantities can be specified as 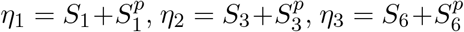 and *η*_4_ = *N* +*N*^*p*^. In order to examine the effect of variation in the amount of STAT1 expression, we numerically construct the bifurcation diagrams with varying *η*_1_ (Fig. 5 (a)). Specifically, for a fixed *η*_1_, the equilibrium points are obtained by numerically finding the solution to **0** = ν ***r***(***x***). To numerically test the stability of each equilibrium point, say ***x***_0_, a small randomly given perturbation *δ****x*** such that (***x***_0_ +*δ****x***) shares the same conserved quantities with ***x***_0_ is exerted to see whether the system state recovers to ***x***_0_.

**Figure 5:**
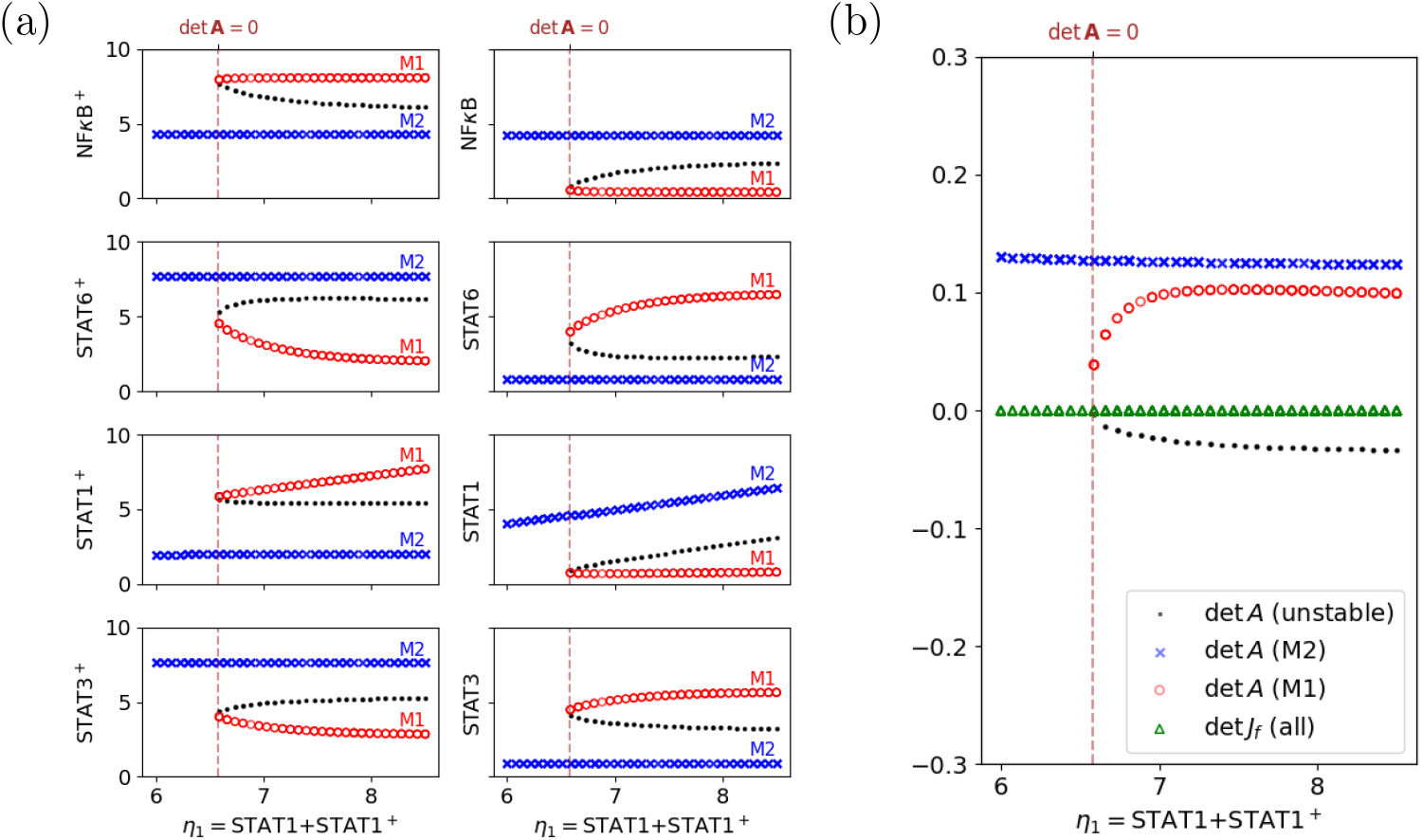
Bifurcation analysis of the WT network. (a) The bifurcation diagrams with the conserved quantity *η*_1_ = STAT1 + STAT1^+^ ranging in [6, 8.5] being the bifurcation parameter. Markers in different shapes are used to denote points on distinct equilibrium branches. The short and long stable equilibrium branches can be interpreted as the M1 (red) and M2 (blue) polarized states, respectively. (b) For each equilibrium as denoted in (a), we numerically estimate the determinants of the matrix ***A*** and the standard Jacobian. It is clear that the positiveness of det ***A*** represents the stability of equilibria, verifying our corollary of stability, and the standard Jacobian determinant always vanishes, failing to be an informative indicator of bifurcation. The vertical dashed line in each of panels illustrates the bifurcation threshold (≈ 6.58) for *η*_1_.

As shown in Fig. 5 (a), the numerical simulations demonstrate that all the chemicals exhibit the bifurcation behavior, which agrees with the SBA prediction. More specifically, there are two stable branches: one with highly-activated NF*κ*B and STAT1, and the other with highly-activated STAT3 and STAT6. Thus, our simple model successfully reproduced the known phenotypes of M1 and M2 macrophages. Besides, one of the stable branches, corresponding to M1 phenotype, disappears when STAT1 expression *η*_1_ decreases to a threshold value (≈ 6.58). This result is qualitatively consistent with the experimental observation that STAT1 deficiency in macrophages leads to a higher ratio of M2 to M1 polarization [23, 24].

Collecting all the stable and unstable equilibrium points in the bifurcation diagram, we shall compute the corresponding det ***A*** and det *J*_*f*_ to see whether the former does outperform det *J*_*f*_ as an indicator for bifurcation and stability. Based on the network topology, we have

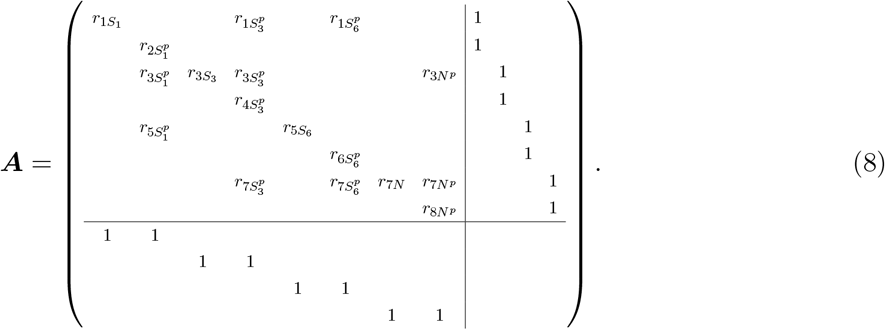

We numerically evaluate det *J*_*f*_ and det ***A*** at all the equilibrium points and confirm that the system is unstable when det ***A*** ≥ 0, as claimed by **Corollary 1** (Fig. 5 (b)). It is shown that the determinant of standard Jacobian always vanishes and thus fails to be a decent bifurcation indicator; on the other hand, det ***A*** stays informative of the occurrence of bifurcation.

#### 2.3.2 Network alterations create buffering structures, resulting in localized bifurcation behaviors

As aforementioned, it has been suggested that the suppression of SOCS3 can cause the hyperactivation of STAT3 and prevent the metastasis of skin cancer [22]. Therefore, one may wish to examine the influence of deleting SOCS3 on the bifurcation behavior associated with macrophage polarization. SOCS3 can be induced by STAT1^+^, STAT3^+^, and NF*κ*B^+^ and negatively regulates STAT3 activation. In the WT network, presented in Fig. 4, these regulations are depicted by the three arrows acting on the activation of STAT3. In the matrix ***A*** corresponding to the WT network, these three regulations are expressed by negative partial derivatives 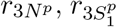, and 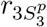 Therefore, the deletion of SOCS3 corresponds to removing these negative regulations from the network and setting these partial derivatives to zero in ***A***. As we will demonstrate below, this network alteration creates buffering structures, which can confine bifurcation behaviors.

We follow the 6-step standard procedure to predict the bifurcation behavior based on the network topology (see Fig. 8 (b) in Appendix B for the altered network). In the first step, the matrix ***A*** is constructed as given in Eq. (8) but with 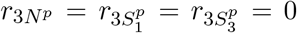, since the regulation effects disappear due to the SOCS3 deletion. In step 2, the matrix ***A*** can be rearranged to be

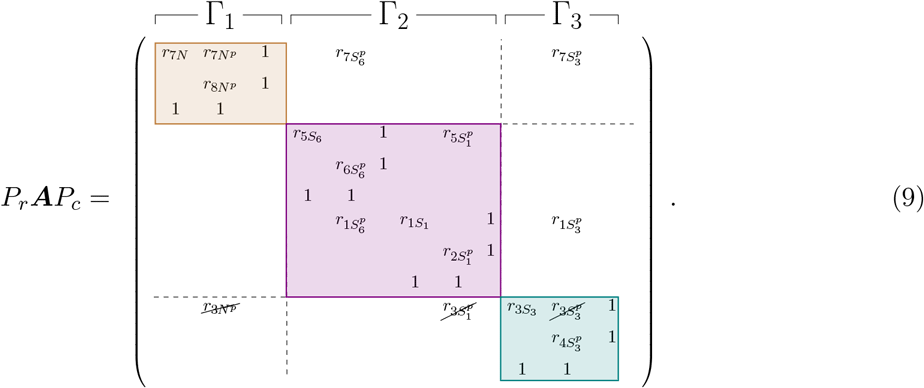

According to the form, we define three subnetworks by

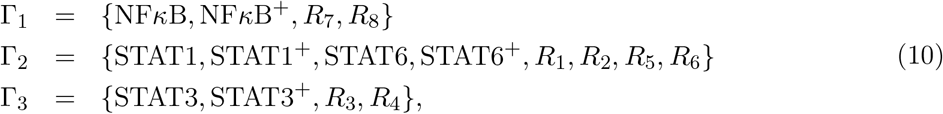

Then, Γ_1_ and (Γ_1_ ∪ Γ_2_) are two buffering structures, and the hierarchy of buffering structures with respect to the inclusion relations can be summarized as step 3 (Fig. 6). Next, calculating the determinants of square blocks in the diagonal in Eq. (9), then one obtains

**Figure 6:**
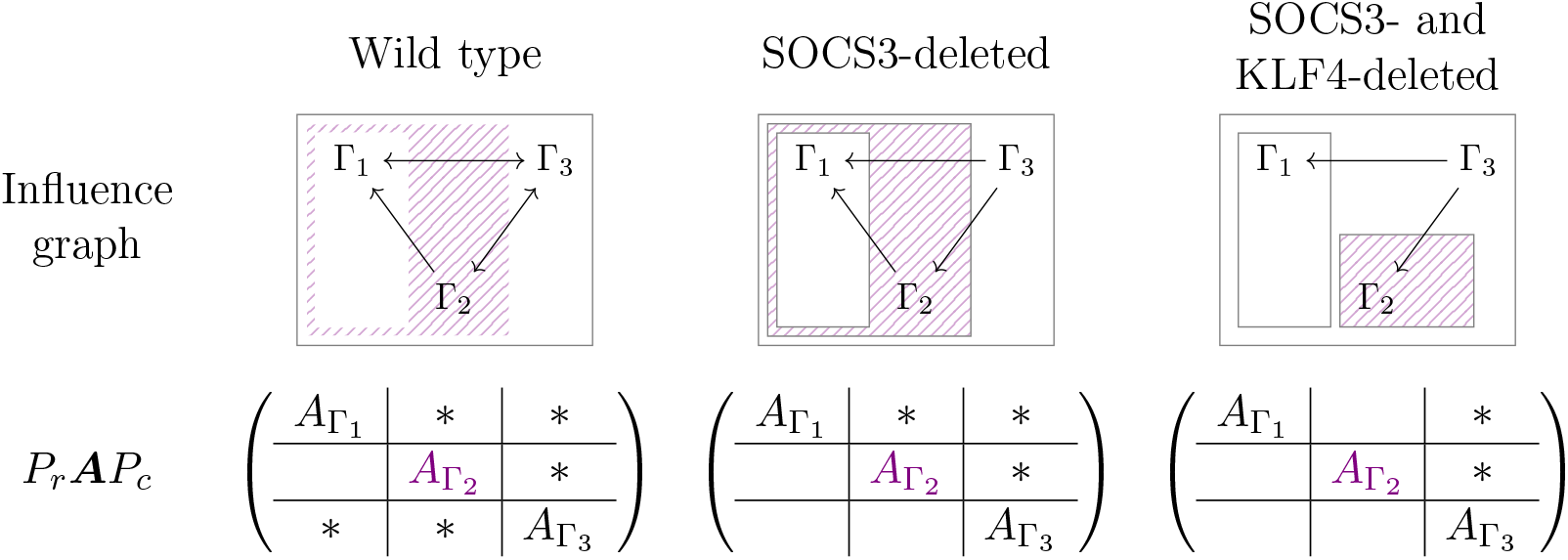
The influence graphs and the rearranged ***A*** matrices corresponding to the wild, the SOCS3-deleted, and the SOCS3-/KLF4-deleted types. In the rearranged matrix, we color the block 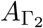 since the sign of its determinant can change, determining the occurrence of bifurcations. With the influence graphs, SBA predicts that chemicals in (Γ_1_ ∪ Γ_2_) exhibit the bifurcation behavior in the SOCS3-deleted network, while the bifurcation behavior is limited within Γ_2_ when KLF4-deletion is further applied to the network.

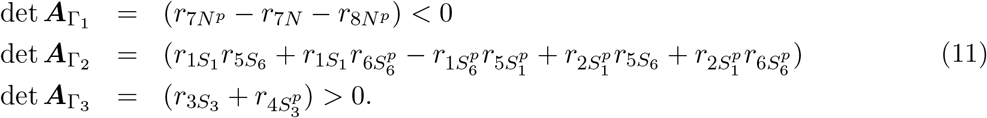

Notice that 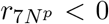 because NF*κ*B^+^ negatively regulates *R*_7_ (i.e., the activation of NF*κ*B), and *r*_7*N*_ *>* 0 and 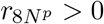 because the NF*κ*B and NF*κ*B^+^ are reactants in *R*_7_ and *R*_8_, respectively. This assures that det 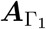 never becomes zero but stays negative, and we analogously have det 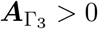 *>* 0. Therefore, as the conclusion of step 5, only Γ_2_ is responsible for the occurrence of bifurcation. Finally, since the minimal buffering structure including Γ_2_ is (Γ_1_ ∪ Γ_2_), the bifurcation behavior can be exhibited only in (Γ_1_ ∪ Γ_2_).

To verify the prediction of SBA that the SOCS3-deleted network can exhibit the bifurcation only in activated and inactivated forms of NF*κ*B, STAT1, and STAT6 (that is, chemicals in (Γ_1_ ∪ Γ_2_)), we adopt the ODE model built for the wild type but with the regulations mediated by SOCS3 removed (see Fig. 8 (b) in Appendix B). The bifurcation diagram numerically computed aligns perfectly with the theoretical prediction based on SBA. Furthermore, in our numerical analysis, the protein STAT3—located outside the buffering structure, (Γ_1_ ∪ Γ_2_), and therefore not exhibiting bifurcation–—persists in a highly activated state, a characteristic akin to M2 polarization. This result is consistent with an experimental report that STAT3 is highly activated in the SOCS3-deleted mutants [22].

In the analysis of the SOCS3-deleted network described above, STAT3 is fixed in the M2-like phenotype due to a buffering structure that confines the bifurcation range within it. To further illustrate our method, we additionally modify this network to fix a subsystem to the M1-like phenotype. Equation (9) suggests that Γ_2_ becomes an independent buffering structure if 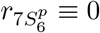 in the SOCS3-deleted network. This partial derivative is corresponded to the regulation of STAT6^+^ on the activation of NF*κ*B via the gene expression of KLF4 (Krüppel-like factor 4). When the deletions of both SOCS3 and KLF4 are applied (see Fig. 8 (c) in Appendix B for the doubly-altered network), we still have Eq. (11); namely, only Γ_2_ is responsible for the occurrence of bifurcation. Moreover, since the minimal buffering structure including Γ_2_ is Γ_2_ itself, SBA concludes that only the four chemicals STAT1, STAT1^+^, STAT6, and STAT6^+^ can exhibit the bifurcation behaviors. With an analogous numerical simulation, we once again verify that the prediction of SBA (Fig. 7 (b)). Only STAT1 and STAT6, located within Γ_2_, exhibit bifurcation, resulting in branches where both of the proteins are M1-like or both are M2-like. Conversely, STAT3 and NF*κ*B, located outside Γ_2_, remain persistently highly activated, each resembling the M2 and M1 phenotypes, respectively. In other words, our results suggest that the double mutations would create a phenotype that simultaneously displays M1- and M2-associated features, robustly (e.g. independently of *η*_1_).

**Figure 7:**
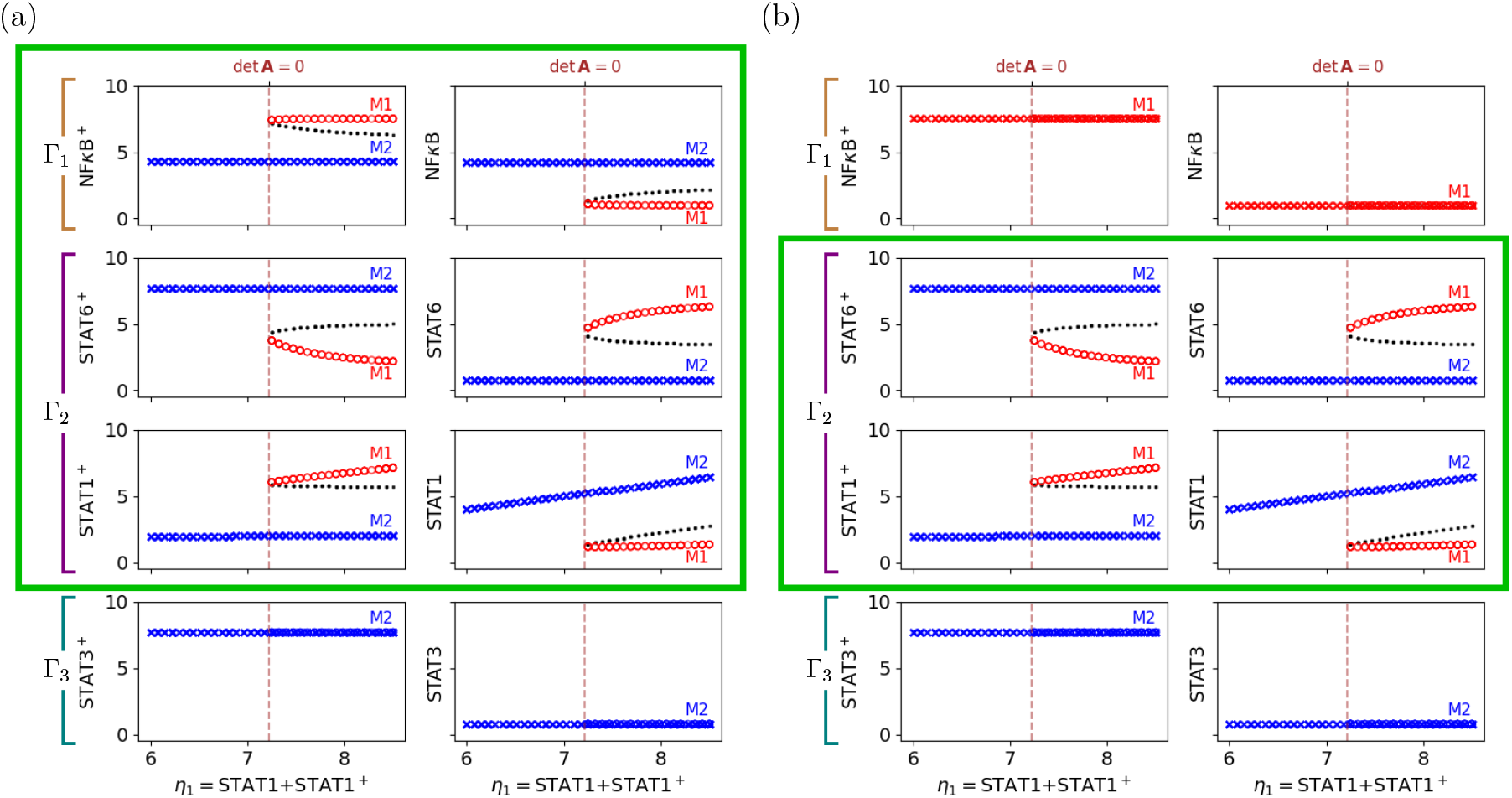
Bifurcation analysis of the modified networks. The bifurcation diagrams of (a) the SOCS3-deleted network and (b) the SOCS3- and KLF4-deleted network, which fully agree with the prediction of SBA. Markers in different shapes denote distinct equilibrium branches as done in Fig. 5, where as the coloring for each chemical is determined by the biological interpretation (red for M1-like, and blue for M2-like). The vertical dashed lines denote the bifurcation threshold (≈ 7.25) for *η*_1_.

**Figure 8:**
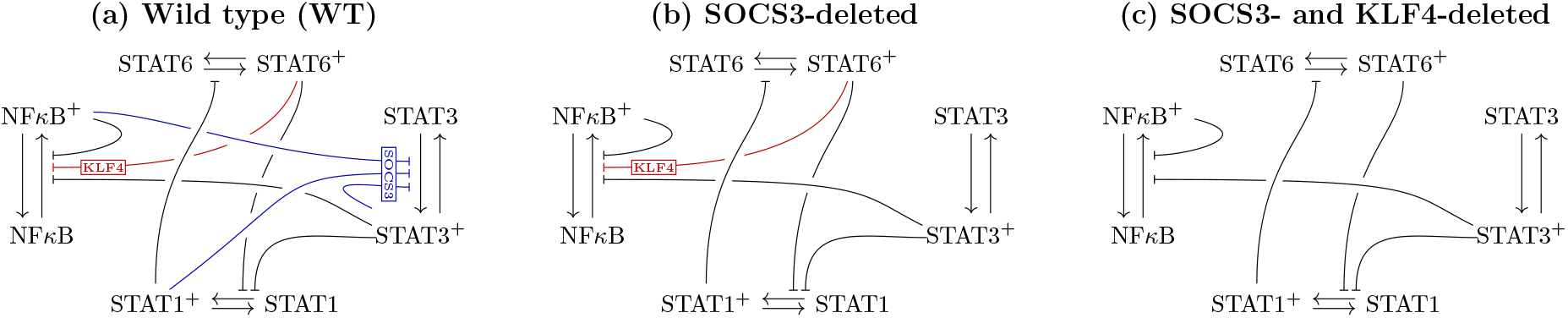
The reaction networks for (a) the wild type (WT), (b) the SOCS3-deleted (altered) mutants, and (c) SOCS3- and KLF4-deleted (dou_1_bly-altered) mutants.

## 3 Discussion

In this paper, we further extend structural bifurcation analysis, which determines the fundamental properties of bifurcation behavior in chemical reaction systems based solely on network structures. We have increased the theoretical generality of the method while also demonstrating its practicality for elucidating biological systems. We have shown, through rigorous mathematical formulae, the direct relationship between the Jacobian matrix, which characterizes conventional theory, and the matrix ***A***, which characterizes structural bifurcation analysis. This not only clarifies the relationship with conventional theory, but also makes it possible to analyze bifurcation behavior in response to changes in conserved quantities in addition to changes in reaction parameters. We also showed the method of explicitly obtaining the influence relationship of parameter perturbation on substructures on the network as an influence graph from the matrix ***A***. Furthermore, we showed the algorithm for comprehensively performing structural bifurcation analysis and structural sensitivity analysis. These results were applied to the macrophage polarization system, and not only did they clarify the mechanisms behind cell fate regulation, but they also enabled us to predict changes in cell fate switching behavior in mutants.

As the example of the macrophage polarization system shows, bifurcation has important significance for biological systems, as it creates discrete states and transitions between them. Nevertheless, bifurcation analysis has not yet achieved sufficient popularity in the life sciences. The main reason for this is the difficulty of the analysis. To find bifurcation phenomena, it is necessary to identify the equilibrium points of the target system and investigate the dependence of the equilibrium points on parameters. To perform these analyses on biological systems, which include many molecular species, i.e., high-dimensional dynamical systems, the practical way is to perform numerical analysis. Nonetheless, such a high dimensional biological system always comes with a large number of parameters involved. Therefore, until now, the only way to identify the equilibrium points and parameter dependencies of high-dimensional dynamical systems in life was through a trial-and-error approach using numerical calculations.

The structural bifurcation analysis we propose has the potential to address the aforementioned issues by determining fundamental bifurcation properties solely based on network structure in a model-free manner. This approach becomes especially powerful when buffering structures, identified according to the network structures, are present. By decomposing large biological systems into substructures determined by buffering structures, we can elucidate the structural conditions necessary for bifurcation, and pinpoint the bifurcation parameters and variables that exhibit bifurcation behavior within each substructure of the network.

The matrix ***A*** directly reflects the topology of the network and provides deeper insight into the network structure for analysis. The new mathematical finding presented in this paper not only generalizes the theory of structural bifurcation analysis to chemical reaction networks with conserved quantities but also opens avenues for further expansion of the theory. For instance, it implies that the (in-)stability of the system can be inferred from the matrix ***A***. We anticipate that this breakthrough can be integrated into other methods rooted in the analyses of Jacobian matrix. For example, the type of bifurcation can be determined by the eigenspectrum of Jacobian matrix [26], and the sign of Jacobian determinant can give a hint whether the system possesses a single equilibrium point [27]. We believe that using the matrix ***A*** instead of Jacobian matrix can render these theories a structural insight into chemical reaction networks.

With the network in Example 1, we obtained three different bifurcation scenarios by assuming three different mathematical models, i.e., sets of reaction rate functions. While different reaction rate functions yielded different bifurcation types, the use of a common network produced consistent results for bifurcation conditions, bifurcation parameters, and bifurcating variables. This clearly demonstrates that these three dynamical properties are determined solely by the structure of the reaction network. Historically, bifurcation patterns have been a significant focus in the mathematical sciences. However, within the context of structural bifurcation analysis, bifurcation patterns may be regarded as incidental properties that emerge only after the network structure has been defined and a specific model is assumed.

The structural bifurcation analysis presented in this study will serve as a powerful tool for advancing our understanding of biological systems. To demonstrate this, we analyzed the macrophage polarization system and derived general predictions derived from the network structures. In wildtype cells, macrophage polarization was examined by varying the expression level of STAT1. When STAT1 expression is low, macrophages are predicted to exhibit only the M2 type (anti-inflammatory), lacking diversity in phenotype. Conversely, when STAT1 expression is high, both M1 (inflammatory) and M2 types appear. In SOCS3 mutants, our analysis indicates that STAT3 keeps highly activated, independent of STAT1 expression, without exhibiting bifurcation behavior. This finding aligns with experimental observations of STAT3 hyperactivation in SOCS3 mutants. However, it is important to note that bifurcation behavior can still occur in the activities of STAT1 and STAT6 in SOCS3 mutants. This result suggests that the polarization behavior of macrophages with SOCS3 deletion may differ depending on which gene’s expression is observed. For instance, STAT3 may always display hyperactivation and an M2 phenotype, while STAT1 and STAT6 show normal polarization into either M1 or M2, similar to the wild type. Additionally, structural bifurcation analysis suggests that the suppression or deletion of KLF4 in the SOCS3-knockout system further eliminates multistability in both NF*κ*B and STAT3, leading to their simultaneous hyperactivation. Interestingly, while NF*κ*B hyperactivation is associated with the induction of pro-inflammatory genes, STAT3 hyperactivation promotes anti-inflammatory responses. This suggests that a macrophage lacking both SOCS3 and KLF4 may exhibit seemingly contradictory behaviors, warranting further investigation to understand its role in inflammation.

We believe these predictions can be verified through experiments that are relatively easy to perform. Furthermore, since these predictions were made model-free and based solely on the network structure, verification experiments could enhance the accuracy of the reaction network information. In other words, if the predicted behaviors are not observed in the experiments, it could indicate that the structure of the reaction network is incorrect. By adjusting the network structure as needed, deriving new predictions, and conducting further experimental verification, we can obtain more accurate information about the reaction network. Finally, it is noteworthy that the M1 and M2 dichotomy is a simplistic and convenient categorization of possible macrophage phenotypes. There are subtypes between the M1 and M2 phenotypes, and their immunological roles are increasingly recognized as important [28, 29]. Our structural method may also be beneficial in predicting potential subtypes based on a network structure or in proposing modifications to achieve targeted phenotypes.

## 4 Method

### 4.1 Review of Structural Bifurcation Analysis (SBA)

Suppose that a network is given by *N* reactions

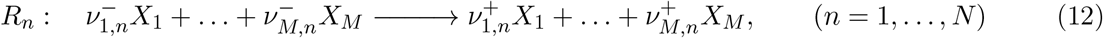

where 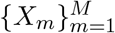 is the collection of chemical species and 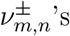 represent the coefficients in the reactions. Then, an *M* × *N* stoichiometric matrix ν is defined by 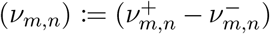, encoding the network topology as a weighted incidence matrix of the directed graph. Then, the dynamics follows the ordinary differential equations

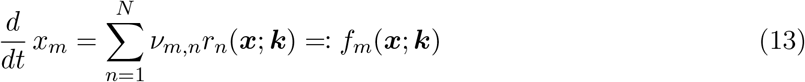

for each *m* = 1, 2, …, *M*, where ***k*** is a parameter vector in the reaction rate function ***r*** : ℝ^*M*^ ×ℝ^*N*^ → ℝ^*N*^. There is no universal rule of determining the functional form the of reaction rate function ***r***. The nonlinearity of reaction rate functions in general contributes to the difficulties of modeling approaches in quantitatively estimating the parameters based on data. The influences of chemicals to reaction rates, meanwhile, can usually be qualitatively determined by the biological background knowledge. For instance, the value of the partial derivative 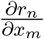 is positive if *X*_*m*_ is a reactant in the reaction *R*_*n*_, and it is negative if *X*_*m*_ negatively regulates the reaction as a suppressor, and it can also be zero when *X*_*m*_ does not affect the reaction rate of *R*_*n*_.

Without functional forms of kinetics, the equilibrium dynamics can still be well characterized to a degree with great helps of two linear subspaces. When the system is at an equilibrium point 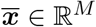, the change rate is zero and thus we have

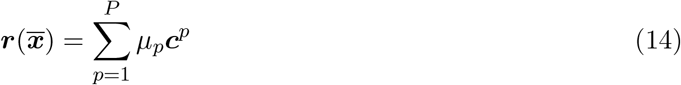

for some *µ* ∈ ℝ^*P*^, where 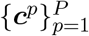 is a basis of ker ν ⊆ ℝ^*N*^ and *P* = dim ker ν. We may put *C* := (***c***^1^, …, ***c***^*P*^) ∈ ℝ^*N ×P*^ and let ⟨*C*⟩ denote the space spanned by column vectors of *C*. Besides, we say a chemical reaction network has a conserved quantity if a linear combination of chemicals is time-invariant, namely, 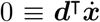 for some ***d*** ∈ ℝ^*M*^ for any ***x*** ∈ ℝ^*M*^. By the dynamics given by Eq. (13), we see ***d*** ∈ ker ν^⊺^ ≃ coker ν. Therefore, take a cokernel basis *D* = (***d***^1^, …, ***d***^*Q*^) ∈ ℝ^*M ×Q*^ with *Q* = dim ker ν^⊺^, then ***η*** := *D*^⊺^***x*** specifies all the conserved quantities. We similarly denote ⟨*D*⟩ = ker ν^⊺^.

With the graph and the biological information rendered by the chemical reactions, a square matrix ***A*** is defined as

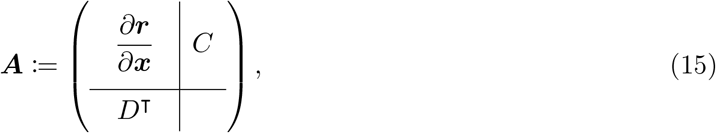

of which lower-right block is empty (throughout the article, trivial blocks are denoted empty). Together with *C* indicating loops of reactions, *D* determining the conserved quantities, the zero-nonzero pattern of entries of the matrix ***A*** directly encodes the structure of the reaction network structure. Such a matrix was first introduced for the sensitivity analysis of chemical reaction networks [30, 31]; specifically, a sensitivity matrix ***S*** = −***A***^−1^ determines the system responses to perturbations in parameters. More explicitly,

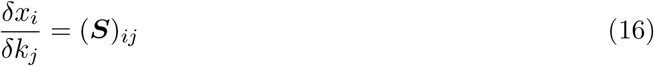

indicates the change (*x*_*i*_ ↦ *x*_*i*_ + *dx*_*i*_) in the chemical *X*_*i*_ in response to a parameter change (*k*_*i*_ ↦ *k*_*i*_ + *dk*_*i*_) in *K*_*j*_ (see Appendix A.I for details of structural sensitivity analysis).

The identification of buffering structures is important in the approach of structural analysis, since a localization principle arises from buffering structures—the influence of a parameter change inside a buffering structure localizes inside the buffering structure (see Appendix A.III.). The localization principle does not hold only in the context of the aforementioned sensitivity problem, but it was extended to bifurcation analysis by the mathematical finding of Okada *et al*. [13, 14] *that det* ***A*** ∝ det *J*_*f*_ when ν is full-ranked (i.e., the system has conserved quantities), where *J*_*f*_ being the Jacobian of *f* in the Eq. (13). This leads to the development of Structural Bifurcation Analysis, in which a bifurcation is identified by the vanishing of the determinant of ***A***. In particular, as shown in Eq. (4), the identification of buffering structures facilitates the factorization of the determinant (followed by the fact that det 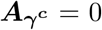). The presence of multiple buffering structures (say *γ*_*α*_’s) highly benefits us in bifurcation analysis with the capability of

1. Factorizing the bifurcation conditions: the bifurcation analysis of a complex network can be facilitated by the network decomposition into determinant structures (denoted by Γ_*β*_’s), which are defined as buffering structures with subtraction of their inner buffering structures (Fig. 1 (3)). For each determinant structure, the condition for bifurcation occurrence can be examined from its network structure.
2. Finding (bifurcation-)inducing parameters: for each determinant structure Γ_*β*_, the bifurcation parameters associated with det 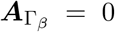, termed as the inducing parameters for Γ_*β*_, are identifiable on the network. The bifurcations are induced by the parameters which are outside any buffering structures non-intersecting with the determinant structure Γ_*β*_ (Fig. 1 (4)).
3. Determining the bifurcating chemicals: for each determinant structure Γ_*β*_, the chemicals exhibiting bifurcation behaviors associated with det 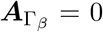, termed as the bifurcating chemicals for Γ_*β*_, are identifiable on the network. When the bifurcation condition is satisfied, the bifurcation of steady-state concentrations (and fluxes) appears only inside the (minimal) buffering structure that contains the determinant structure Γ_*β*_ (Fig. 1 (5)).

### 4.2 The generalization of SBA

SBA is rooted in the finding of Okada *et al*. that det ***A*** ∝ det *J*_*f*_, and we remark that the theorem is proved under the assumption that ν is full-ranked, that is, no conserved quantity is present in the considered network [13, 14]. It remains an open problem whether the framework of SBA can be employed when conserved quantities exist.

What is the problem with the conserved quantities? When the cokernel space ⟨*D*⟩ is nontrivial, we have 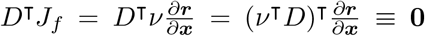. This implies that the Jacobian *J*_*f*_ is always noninvertible because of the vanishing eigenvalues due to the nontrivial cokernel space. Intuitively speaking, a trajectory ***x***(*t*) always lies on a hyperplane parallel to im ν, which does not span the whole space when ν is not full-ranked. Therefore, *L* of the eigenvalues that measure diverging rates of trajectories must be zero, where *L* := dim coker ν = (*M* −dim im ν) is the number of independent conserved quantities.

To circumvent the problem with the redundant eigenvalues, the intuition above motivates us to have a change of coordinates with (im ν) and its orthogonal complement (im ν)^⊥^ being the two axes. To do so, we put *V* to be a full-ranked matrix such that ⟨*V* ⟩ = im ν and consider the pseudo-inverses

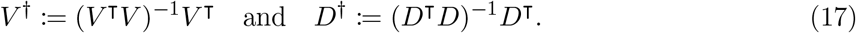

Then, a linear isomorphism for the desired change of coordinate is obtained as

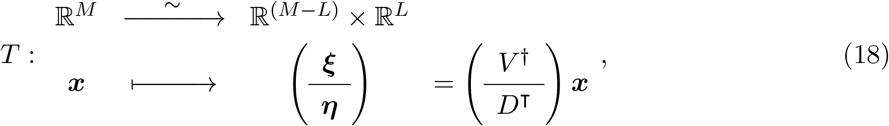

or, alternatively, ***x*** = *V* ***ξ*** + (*D*^*†*^)^⊺^***η***. Then, the dynamics can be rewritten as

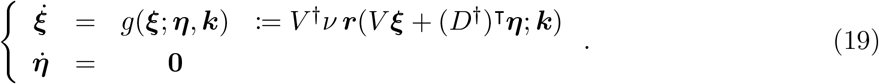

This gives a clear picture showing that the dynamics of ***x*** solely counts on that of ***ξ***, and thus it is sufficient to consider the Jacobian *J*_*g*_ = *V* ^*†*^*J*_*f*_ *V* of *g* for the bifurcation and stability analyses. By considering *J*_*g*_ instead of *J*_*f*_, the problem of the vanishing eigenvalues due to conserved quantities is well circumvented. Moreover, the Jacobian *J*_*g*_ is associated with the matrix ***A*** which is given by the network topology. Considering a constant matrix

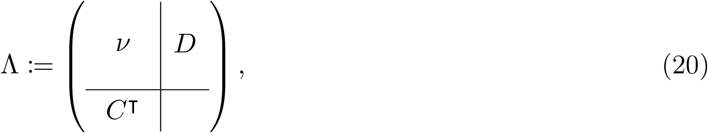

we have the relation associating ***A*** and Jacobian, given by

#### Theorem 1

*Given a kernel basis C and a cokernel basis D of the mapping* ν, *then there exists an orthogonal basis* 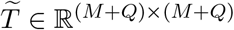 *such that*

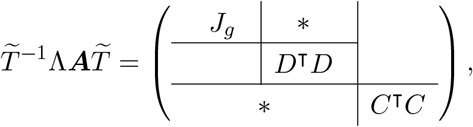

*and hence* det 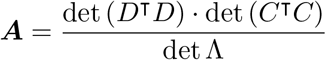 det *J*.

*In particular, when* dim coker ν = 0, *we have* det 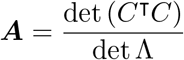 det *J*.

The relation further gives **Corollary 1** as a criterion for stable equilibria. The proofs are left in Appendix C: The theorems and the proofs.

### 4.3 An example network with conserved quantities

To show the problem induced by the presence of conserved quantities and how our theorem circumvent it, we consider a simple network.

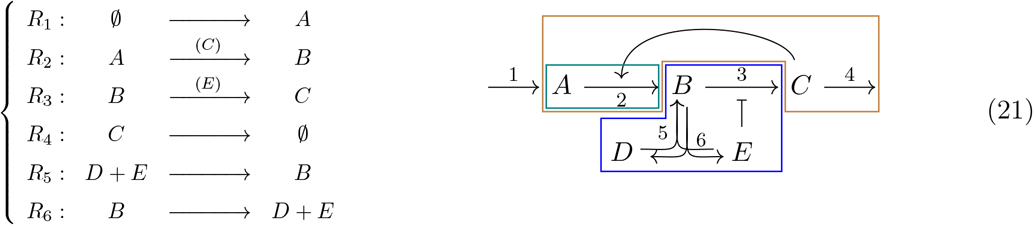

The corresponding stoichiometric matrix is

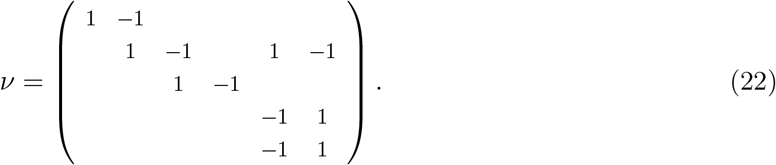

Furthermore, one sees that the cokernel space is given by

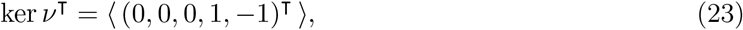

since *η*_1_ := 1 · *x*_*D*_ + (−1) · *x*_*E*_ determines the conserved quantity. On the other hand, the kernel basis can be obtained by considering the two loops 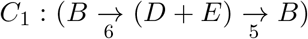 and 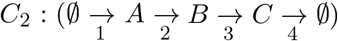 As such, one obtains

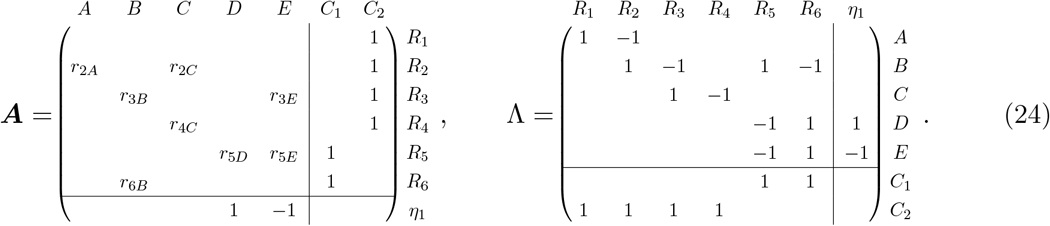

By the identification as given in Appendix A: Buffering structures and the localization principle, it can be seen that Γ_1_, (Γ_1_ ∪ Γ_2_) and Γ_3_ are three buffering structures, where Γ_1_ := {*A, R*_2_}, Γ_2_ := {*C, R*_4_}, Γ_3_ := {*B, D, E, R*_3_, *R*_5_, *R*_6_}, and Γ_4_ := {*R*_1_}. We have the rearranged form

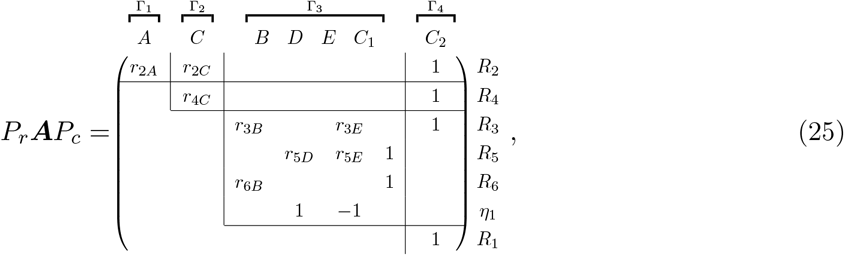

where *P*_*r*_ and *P*_*c*_ are permutation matrices with det *P*_*r*_ = det *P*_*c*_ = −1. The step of rearranging ***A*** according to the buffering structure highly facilitates the computation and factorization of the determinants, giving

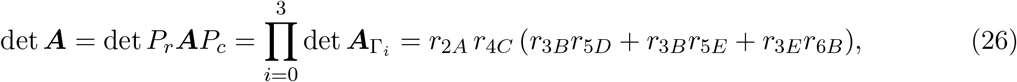

in which 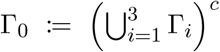 is the complement of the union of all buffering structures. With **Theorem 1**, it is known that det 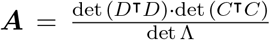·det *J*_*g*_ = det *J*_*g*_ since det (*C*^⊺^*C*) = 8, det (*D*^⊺^*D*) = 2, and det Λ = 16; meanwhile,

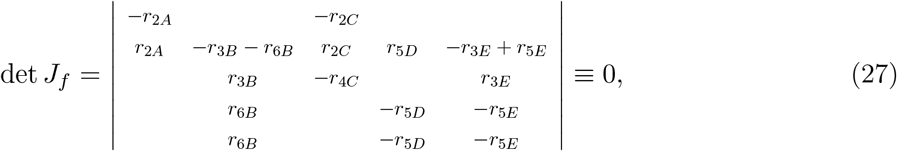

which always vanishes and thus fails to be an informative bifurcation indicator.

Despite the vanishing standard Jacobian determinant, we see that the generalized SBA can be applied to this network to predict the bifurcation behaviors. Since *A* is a reactant in *R*_2_ and *C* is a reactant in *R*_4_, both *r*_2*A*_ and *r*_4*C*_ do not vanish but stay positive. The Eq. (26) implies that a bifurcation occur only when det 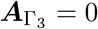. Furthermore, Γ_3_ is a buffering structure, one can conclude that the bifurcation behaviors can be exhibited only by chemicals *B, D*, and *E*.

### 4.4 Influence graph among subnetworks

As shown by the two pedagogical examples, a network decomposition into subnetworks readily follows the rearrangement of the matrix ***A***. Explicitly, the square blocks in the diagonal of the upper block triangular *P*_*r*_***A****P*_*c*_ define disjoint subnetworks Γ_*i*_’s. More specifically, the rearranged matrix *P*_*r*_***A****P*_*c*_ takes the form

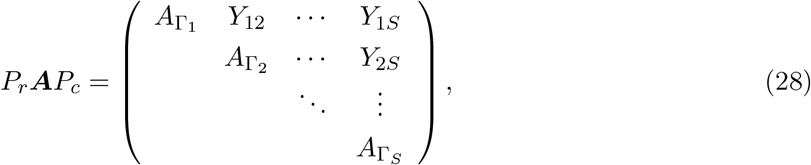

and the subnetworks Γ_*i*_’s can be determined accordingly: a subnetwork Γ_*i*_ consists of a chemical/reaction if a Γ_*i*_’s column/row is corresponding to the chemical/reaction. We term the subnetworks Γ_*α*_’s the determinant structures, which are not necessarily buffering structures, but a buffering structure is always a union of some determinant structures. In other words, determinant structures give a finer network decomposition then buffering structures do.

Our interest is the *influence* relation among these subnetworks Γ_*i*_’s, which is termed determinant structures. Before introducing how the relation is revealed by SBA, we shall first clarify what it means by *influence*. We say a subnetwork Γ_*α*_ influence another subnetwork Γ_*β*_ if there exists some parameter *K*_*i*_ ∈ Γ_*α*_ and *X*_*j*_ ∈ Γ_*β*_ such that 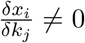, of which idea is identical to Eq. (16).

The influence relation among the determinant structures is encoded by blocks *Y*_*ij*_’s in the matrix representation (28). In particular, a subnetwork Γ_*j*_ influences another subnetwork Γ_*i*_ if *Y*_*ij*_ is not a zero matrix. As such, we can draw a directed graph to visualize the influencing relation among subnetworks, by considering the adjacency matrix *G* [32] defined by

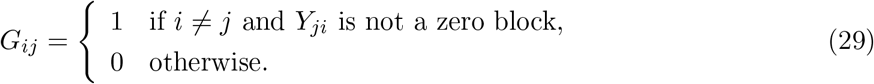

We may take the network given in Eq. (21) for example, with Γ_1_ := {*A, R*_2_}, Γ_2_ := {*C, R*_4_}, Γ_3_ := {*B, D, E, R*_3_, *R*_5_, *R*_6_}, and Γ_4_ := {*R*_1_}, according to the rearranged form *P*_*r*_***A****P*_*c*_ as in Eq. (25). The matrix can be rewritten into the form

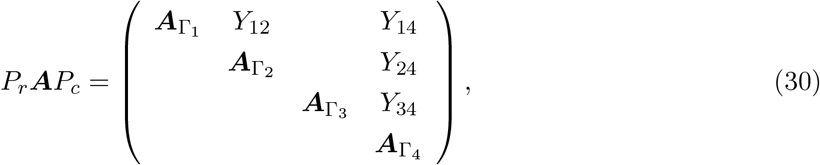

suggesting the adjacency matrix *G* of the influence graph as

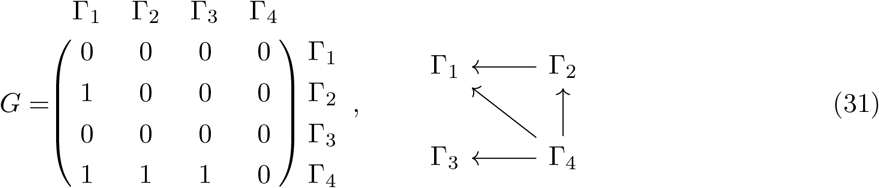

so that the influence relation among determinant structures can be clearly visualized. A determinant structure Γ_*i*_ influences another subnetwork Γ_*j*_ if there is a directed path from Γ_*i*_ to Γ_*j*_ in the influece graph. With the graph, it is straightforward to see that Γ_1_, (Γ_1_ ∪Γ_2_) and Γ_3_ are buffering structures, since they do not have their downstreams in the influence graph. It is clear to see that parameter changes in Γ_2_ could possibly influence Γ_1_ but not Γ_3_ and Γ_4_, since that only Γ_1_ is in the downstream for Γ_2_. It is also worthy of mention that the graph is acyclic (i.e., with no directed cycles), which means that a subnetwork would not be simultaneously in the upstream and the downstream for any other subnetwork. In summary, the influence graph renders the network a hierarchy of subnetworks, giving a structural insight into the equilibrium dynamics.

## Acknowledgements

This research was supported by the CREST program (grant no. JPMJCR1922, JPMJCR24Q4) of the Japan Science and Technology Agency (JST) (http://www.jst.go.jp/EN/index.html), Grant-in-Aid for Scientific Research on Innovative Areas (grant no. 19H05670), Joint Usage/Research Center program of Institute for Life and Medical Sciences Kyoto University. TO was supported by JSPS KAKENHI (Grant No. JP22K03453, JP22K06347) and the RIKEN iTHEMS Program.

## Appendix A: Buffering structures and the localization principle

In this appendix, we are going to formally define *buffering structures* of chemical reaction networks and to explain the localization principle arising from buffering structures. The content is virtually a summary of previous studies [15, 13, 14] to make the present paper self-contained. To help readers better follow up the theory, we will first introduce (i) the theory of Structural Sensitivity Analysis, give (ii) the definition of buffering structures, and finally state and prove (iii) localization principle arising from buffering structures.

### I. Structural Sensitivity Analysis

We shall first start from the ordinary differential equation

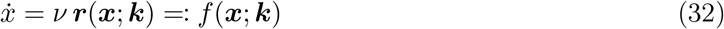

as given in the beginning of Sec. 2.1 in the main text. We may assume the parameter ***k*** in the reaction rate function *f* (***x***; ***k***) = ν ***r***(***x***; ***k***) is a vector in ℝ^*N*^, representing enzyme activities or any other reaction-specific factors. The specificity infers that 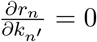 if and only if *n* = *n*^*′*^.

Suppose that the equilibrium manifold is parameterized by the parameter ***k*** and the conserved quantity ***η*** := −*D*^⊺^***x***, locally near a fixed equilibrium point ***x***_0_ ∈ ℝ^*M*^. Besides, put *C* to be a full-ranked matrix such that ⟨−*C*⟩ = ker ν. Remark that we multiply the column vectors of kernel basis matrix and cokernel basis matrix (that is, *C* and *D*) just for simplicity of the following computation. With the parametrization of equilibria, we have

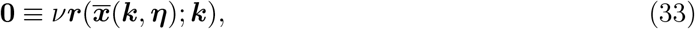

which is followed by the existence of a mapping ***µ***(***k, η***) ∈ ℝ^dim ker ν^ locally such that

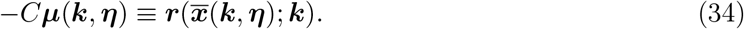

Taking the partial derivative with respect to ***k***, one obtains

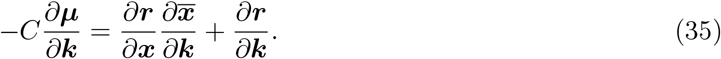

Notice that due to the (enzyme) specificity, the matrix 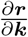 is diagonal; moreover, we may assume that 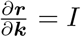 at ***x***_0_, since the unit are flexible to re-scale locally. As such, the equation is rewritten to be

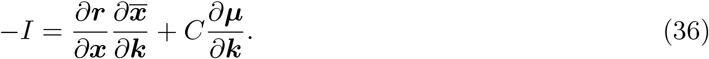

Besides, with 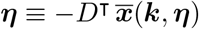 by definition, we have

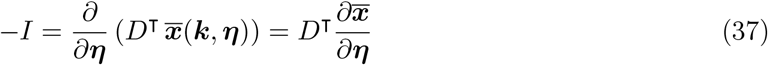

and

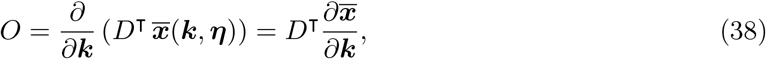

where *O* denotes a zero matrix. Taken the partial derivatives by ***η***, Eq. (34) implies

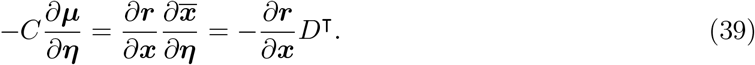

Combine equations (36), (37), (38), and (39), we have

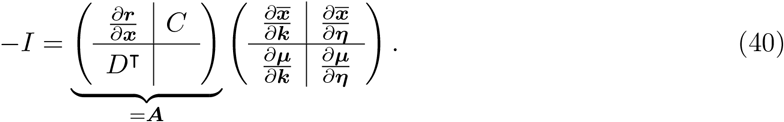

Then, when the matrix ***A*** is invertible at ***x***_0_, the equation (40) implies that the system response (*d****x***, *δ****µ***) to infinitesimal perturbation (*δ****k***, *δ****η***) is given by

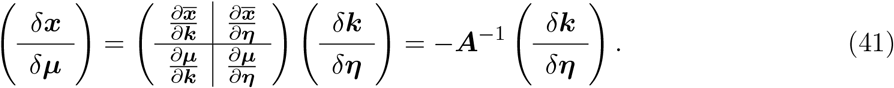

We remark that *δ****x*** and *δ****µ*** respectively represent the changes in the chemical concentrations and the reaction rates (since ***r*** = −*C****µ***).

### II. Buffering structures in chemical reaction network

Next, we are going to provide two ways of defining buffering structures in a chemical reaction network. The two definitions are equivalent when the chemical reaction network is *regular*, that is, det ***A*** ≢0. It is reasonable to consider regular networks only; after all, when a network is not regular, then the system is unstable everywhere in the state space (by **Theorem 1**), which is unrealistic in cell biology.

The first definition is based on an index that depends on the network topology, which highly facilitates the identification of buffering structures when it is done manually. The second definition is based on the matrix representation of ***A***. It comes in handy in mathematical derivations as well as algorithms that identify buffering structures, whereas it requires the kernel basis and cokernel basis (that is, matrices *C* and *D* in Eq. (15)) to be properly chosen.

Adopting the settings in the Method section, we suppose that the dynamics of a chemical reaction network follows equation (13). Let *γ* = *X*_*γ*_ ∪ *R*_*γ*_ be a subnetwork of the considered network, where *X*_*γ*_ and *R*_*γ*_ respectively denote the collections of chemicals and reactions in the subnetwork *γ*. Let ***x***_1_ and ***x***_2_ be vectors which denote the concentrations of chemicals in and out of *X*_*γ*_, respectively, and without loss of generality we may assume the chemicals to be ordered such that ***x*** = (***x***_1_, ***x***_2_). Similarly, let ***r***_1_(***x***) and ***r***_2_(***x***) be the vectors of reaction rates in and out of *R*_*γ*_. Then, the dynamics as described by Eq. (13) is rewritten into

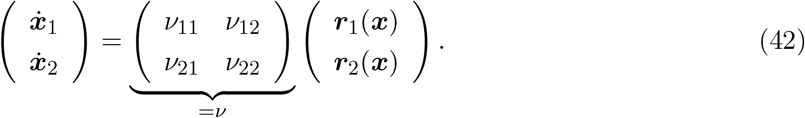

Furthermore, we put

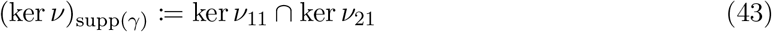

and

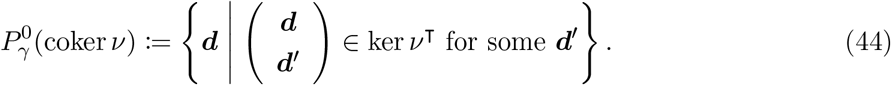

Then, |(ker ν)_supp(*γ*)_| is the number of closed paths within the subnetwork *γ*, and 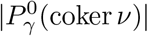 is the number of conserved quantities involved with the subnetwork *γ*. We then define an index by

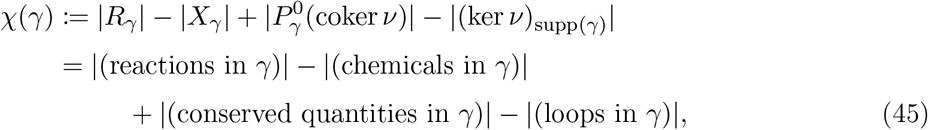

where |*S*| denotes the dimension when *S* is a linear space, and it denotes the cardinal number when *S* is a finite set here. With the index, have the definition

#### Definition 1

*A subnetwork γ is a buffering structure if it satisfies the following two properties:*

1. *(output-completeness) All the reactions regulated by any chemicals of γ are included in γ; namely* 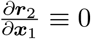
2. *(zero index) The subnetwork satisfies χ*(*γ*) = 0.

or, equivalently,

#### Definition 2

*A subnetwork γ is a buffering structure if there exist full-ranked matrices C and D with* ⟨*C*⟩ = ker ν *and* ⟨*D*⟩ = ker ν^⊺^ *and permutation matrices P*_*r*_, *P*_*c*_ *such that*

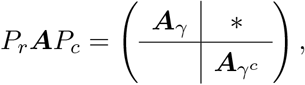

*where* ***A***_*γ*_ *is a square block and γ*^*c*^ *is the complement of γ*.

To see the equivalence, we may let *γ* be an arbitrarily given subnetwork, and let *C*_11_ and *D*_11_ be two matrices with independent column vectors such that ⟨*C*_11_⟩ = (ker ν)_supp(*γ*)_ and 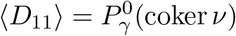 Then, there exist a kernel basis *C* and a cokernel basis *D* in the forms

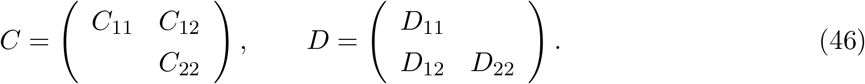

With row- and column-rearrangement, we have the matrix ***A*** in the form

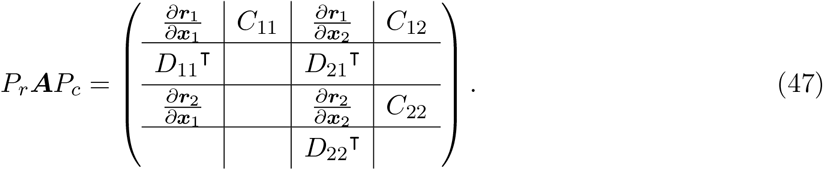

Put

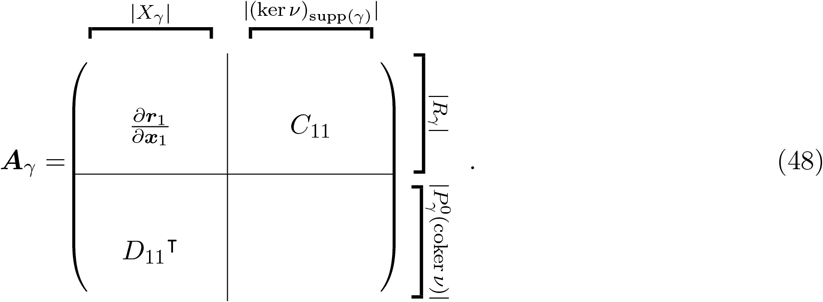

With Eq. (47), we see that *γ* is output-complete if and only if

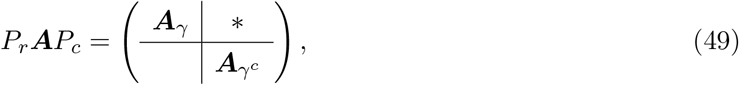

and Eq. (48) shows that ***A***_*γ*_ is a square block if and only if *χ*(*γ*) = 0. This shows the equivalence. Recently, some packages have been released for the identification of buffering structures on Python [33]. The identification of buffering structures is hence feasible even for large networks in pathway databases, in which a pathway map can include hundreds of metabolites [34].

### III. Localization principle arising from buffering structures

Buffering structures are known for confining system responses to parameter changes within itself in two senses. When the system is not at a bifurcation point, the equilibrium dynamics follows the *law of localization* [30, 16]—parameter *changes in a buffering structure do not affect the chemicals outside the buffering structure. When the system bifurcates in response to some bifurcation parameter, then we have the localization principle for bifurcation* [13, 14], *stating that the equilibrium manifold bifurcates only in the chemicals (axes) of the buffering structure γ* only when the corresponding square block ***A***_*γ*_ becomes singular. In this subsection, we clarify the two theorems. The proofs are adopted from the references above but condensed and with notations in the present papers just for readers.

#### A. Law of localization in structural sensitivity analysis

When the matrix ***A*** is invertible at ***x***_0_, the Eq. (40) implies that the system response (*δ*_***x***_, *δ*_***µ***_) to infinitesimal perturbation (*δ*_***k***_, *δ*_***η***_) is given by

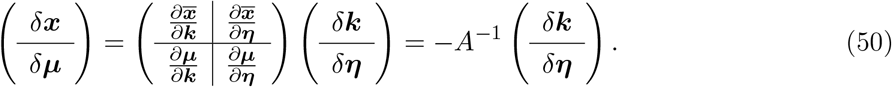

We kindly remark that *δ****x*** and *δ****µ*** respectively represent the changes in the chemical concentrations and the reaction rates (since ***r*** = −*C****µ***). Now, suppose that there is a buffering structure *γ*, then we have matrices *P*_*r*_ and *P*_*c*_ such that

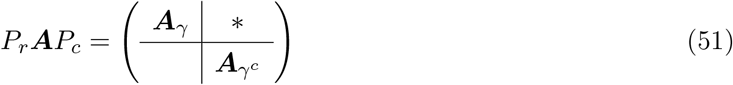

with ***A***_*γ*_ ∈ ℝ^*W ×W*^, where 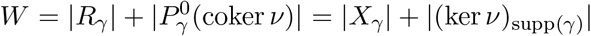 The constructions of the permutation matrices can be given according to the next paragraph.

Adopting the settings of equation (42), we let ***x*** = (***x***_1_, ***x***_2_) denote the concentration of chemicals in and out of a considered buffering structure *γ*. The parameters can be classified as ***k*** = (***k***_1_, ***k***_2_) according to whether the corresp ndin reactions are in the buffering structure or not. Also, we can put ***η*** = (***η***_1_, ***η***_2_) such that 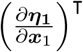 is full-ranked and 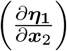 is a zero matrix. Finally, we have ***µ*** = (***µ***_1_, ***µ***_2_) such that 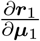 is full-ranked and 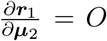 Then, the matrices *P*_*r*_ and *P*_*c*_ inequation (51) can be constructed according to the permutations below:

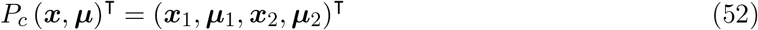

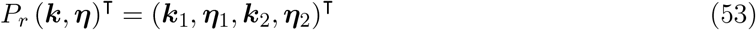

By equation (51) and the assumption that ***A*** is invertible at ***x***_0_, it is seen that

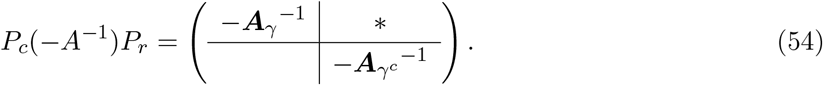

Combine it with equations (40), (52) and (53), then it is shown that

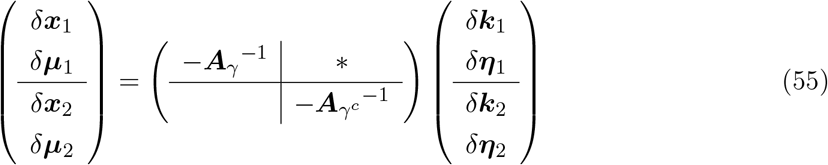

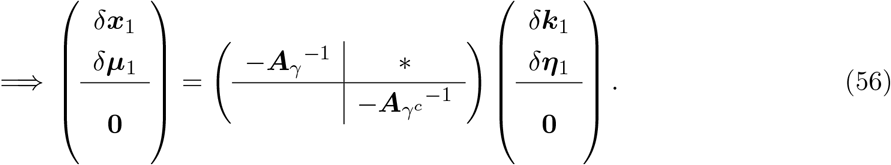

Thus, when the perturbations are only given to parameters and/or conserved quantities of the buffering structure (i.e., *δ****k***_2_ = 0 and *δ****η***_2_ = 0), it does not exert an influence to the outside of the buffering structure (i.e., *δ****x***_2_ = 0 and *δ****µ***_2_ = 0). This proves the law of localization arising from buffering structures in sensitivity analysis. On the other hand, when the perturbation is given to the complement of the buffering structure (namely, either of *δ****k***_2_ and *δ****η***_2_ is non-trivial), it in general influences the buffering structure (that is, *δ****x***_1_ and *δ****µ***_1_).

#### B. Localization principle for bifurcation

We shall first stress that we focus on bifurcations of equilibria, such as transcritical, saddle-node, and pitchfork bifurcations. Bifurcations such as Hopf bifurcation are not considered in the following argument since the interest of the study is the equilibrium dynamics. Therefore, in the neighborhood of a bifurcation point ***x***_0_ ∈ ℝ^*M*^, we may assume that there is a piecewise 𝒞^2^ bifurcation curve

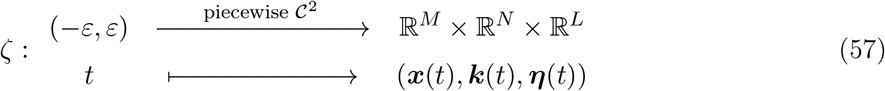

such that ζ(0) = (***x***_0_, ***k***_0_, ***η***_0_) and that 𝒞^2^ and that ***A***|_***x***(*t*)_ is invertible for all *t* ∈ (−*ε, ε*)\{0}.

Once again, we set ***x***(*t*) = (***x***_1_(*t*), ***x***_2_(*t*)), ***k***(*t*) = (***k***_1_(*t*), ***k***_2_(*t*)), and ***η***(*t*) = (***η***_1_(*t*), ***η***_2_(*t*)) accordingly with respect to a considered buffering structure. Suppose that the changes in the parameters and conserved quantities are within the buffering structure, that is, ***k***_2_(*t*) and ***η***_2_(*t*) are constant. Our goal is to show that ***x***_2_(*t*) is a constant as well.

Since ***A***|_***x***(*t*)_ is invertible for all *t* ∈ (−*ε, ε*)\{0}, we see that the law of localization holds over ζ((−*ε, ε*)\{0}); morerover, by the continuity of the bifurcation curve, we see that ***x***_2_(*t*) is constant. In summary, when parameter changes and conserved quantity changes are made within a buffering structure, then the bifurcation behaviors are limited within the buffering structure.

## Appendix B: the models used for method demonstration

### I. A minimal example showing different types of bifurcation

For the chemical reaction network given in the Result section in the main text, we consider three different settings for the kinetics:

**Model (a)**:

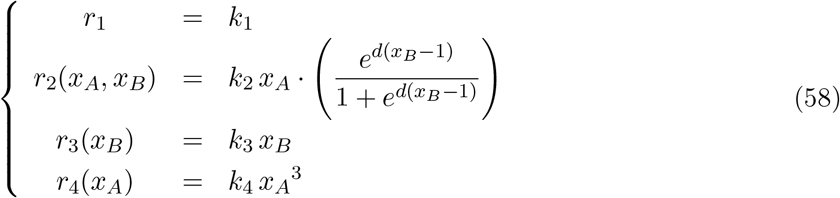

**Model (b)**:

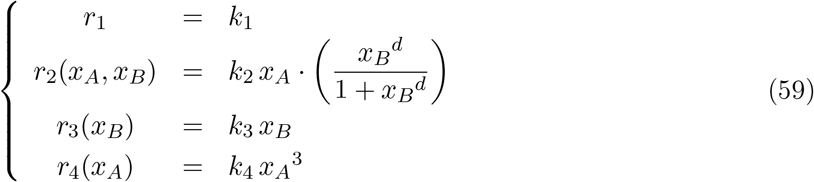

**Model (c)**:

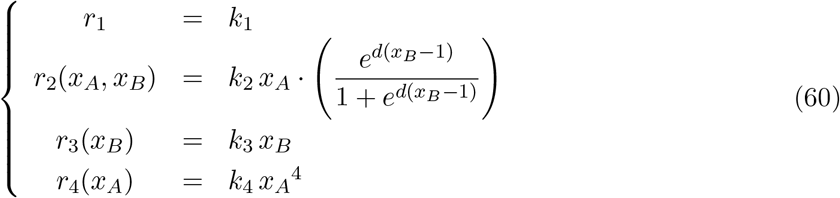

In our numerical simulations, we set *k*_1_ = 8, *k*_2_ = *k*_3_ = *k*_4_ = 1. After searching for bifurcations point for *δ* ∈ [1, 5], we plot the bifurcation diagrams while centering the bifurcation threshold (Fig. 2).

The difference in Model (a) and Model (b) is the sigmoid function that characterizes the regulation of *R*_2_ by *X*_2_, which is the feedback effect of *X*_2_ to itself. On the other hand, Model (a) and Model (c) differs in the form of the rate function *r*_4_. Both in Model (a) and Model (c), *r*_4_ takes a form of the mass-action kinetics but with different powers for the variable *x*_*A*_.

### II. A mathematical model responsible for macrophage polarizarization

In the main text, we employed our method to a network involving parts of JAK-STAT and KF*κ*B signalings. With notations in the main text, for simplicity, we denote the system variables by 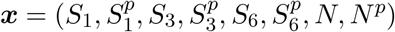 Then, according to the network map, the system follows

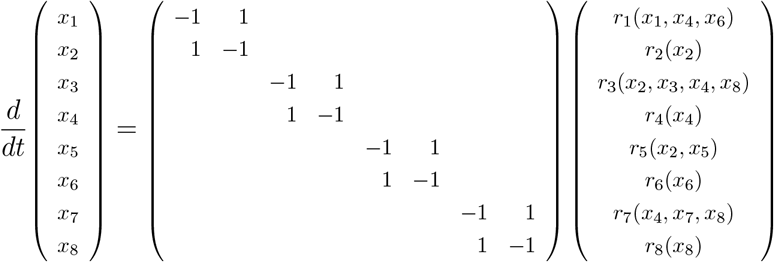

For *r*_*n*_ with *n* being an even number, the reaction is a deactivation, and we may simply put the reaction rate function as

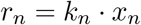

On the other hand, the activation as well as the negative regulation involve phosphorylation, which is commonly modeled as a sigmoid function [25]. Therefore, for *n* = 1, 3, 5, 7, we put

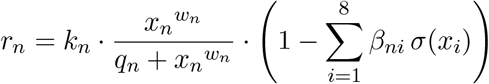

with 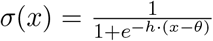, where *β*_*ni*_’s are non-negative numbers such that Σ _*i*_ *β*_*ni*_ ≥ 1. The value of *β*_*ni*_ indicates the strength of regulating effect on *R*_*n*_ by *X*_*i*_; in other words, *β*_*ni*_ *>* 0 if and only if *X*_*i*_ negatively regulate the activation *R*_*n*_ of *X*_*n*_ (*n* = 1, 3, 5, 7, while *i* = 1, 2, …, 8). In the main text, we consider three different scenarios for the networks: (a) the wild type, (b) with SOCS3 deletion, and (c) with SOCS3 and KLF4 deletion (Fig. 8), and we simulate the three types by adjusting the values of *β*_*ni*_’s (Table 1).

**Table 1:** Parameter settings for the numerical experiments of networks responsible for macrophage polarization.

| Parameters |  |  |  |  |  |  |  |  |  |  |  |  |  |  |
| --- | --- | --- | --- | --- | --- | --- | --- | --- | --- | --- | --- | --- | --- | --- |
| $k_1 = k_2 = k_3 = k_4 = k_5 = k_6 = k_7 = k_8 = 1,$ | | | | | $h = 5,$ | | | | | | | | | |
| $q_1 = q_3 = q_5 = q_7 = 1,$ | | | | | $\theta = 0.15,$ | | | | | | | | | |
| $(w_1, w_3, w_5, w_7) = (2, 2, 2, 1),$ | | | | | | | | | | | | | | |
| (a) Wild type |  |  |  |  | (b) SOCS3-deleted |  |  |  |  | (c) SOCS3- and KLF4-deleted |  |  |  |  |
| $\beta = \begin{pmatrix} 0 & 0 & 0 & 0.45 & 0 & 0.45 & 0 & 0 \\ 0 & 0 & 0 & 0 & 0 & 0 & 0 & 0 \\ 0 & 0.45 & 0 & 0.045 & 0 & 0 & 0 & 0.405 \\ 0 & 0 & 0 & 0 & 0 & 0 & 0 & 0 \\ 0 & 0.9 & 0 & 0 & 0 & 0 & 0 & 0 \\ 0 & 0 & 0 & 0 & 0 & 0 & 0 & 0 \\ 0 & 0 & 0 & 0.405 & 0 & 0.405 & 0 & 0.09 \\ 0 & 0 & 0 & 0 & 0 & 0 & 0 & 0 \end{pmatrix}$ | | | | | $\beta = \begin{pmatrix} 0 & 0 & 0 & 0.45 & 0 & 0.45 & 0 & 0 \\ 0 & 0 & 0 & 0 & 0 & 0 & 0 & 0 \\ 0 & 0 & 0 & 0 & 0 & 0 & 0 & 0 \\ 0 & 0 & 0 & 0 & 0 & 0 & 0 & 0 \\ 0 & 0.9 & 0 & 0 & 0 & 0 & 0 & 0 \\ 0 & 0 & 0 & 0 & 0 & 0 & 0 & 0 \\ 0 & 0 & 0 & 0.405 & 0 & 0.405 & 0 & 0.09 \\ 0 & 0 & 0 & 0 & 0 & 0 & 0 & 0 \end{pmatrix}$ | | | | | $\beta = \begin{pmatrix} 0 & 0 & 0 & 0.45 & 0 & 0.45 & 0 & 0 \\ 0 & 0 & 0 & 0 & 0 & 0 & 0 & 0 \\ 0 & 0 & 0 & 0 & 0 & 0 & 0 & 0 \\ 0 & 0 & 0 & 0 & 0 & 0 & 0 & 0 \\ 0 & 0.9 & 0 & 0 & 0 & 0 & 0 & 0 \\ 0 & 0 & 0 & 0 & 0 & 0 & 0 & 0 \\ 0 & 0 & 0 & 0.405 & 0 & 0 & 0 & 0.09 \\ 0 & 0 & 0 & 0 & 0 & 0 & 0 & 0 \end{pmatrix}$ | | | | |

## Appendix C: The theorems and the proofs

The notations are adopted from the main text.

### Theorem 1

*Given a kernel basis C and a cokernel basis D of the mapping* ν, *then there exists an orthogonal basis* 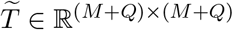 *such that*

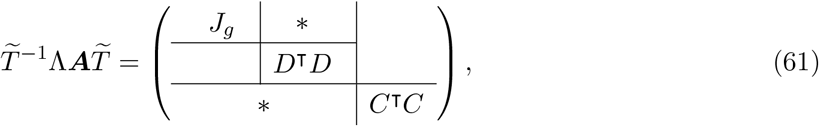

*and hence* det 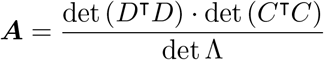. det *J*_*g*_.

*In particular, when* dim coker ν = 0, *we have* det 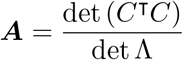·det *J*_*f*_.

(proof) With the fact ν*C* = **0** following ⟨*C*⟩ = ker ν, it is straightforward to see that

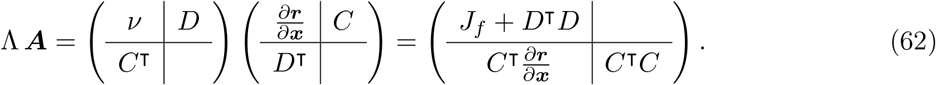

Since *D*^⊺^*V* = 0, *D*^⊺^*J*_*f*_ = 0, and *D*^⊺^(*D*^*†*^)^⊺^ = *I*, we have

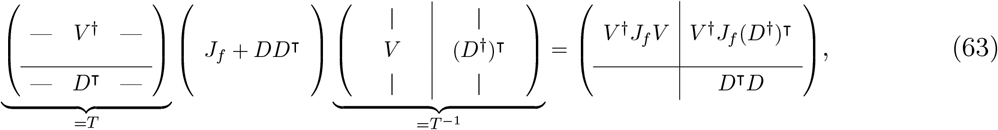

where *T* is the matrix representation of the mapping as given in (18). Put

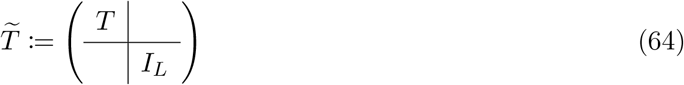

in which *I*_*L*_ is the *L* × *L* identity matrix with *L* = dim coker ν. Then, together with Eq. (62) and Eq. (63) as well as the definition *J*_*g*_ = *V* ^*†*^*J*_*f*_ *V*, we obtain

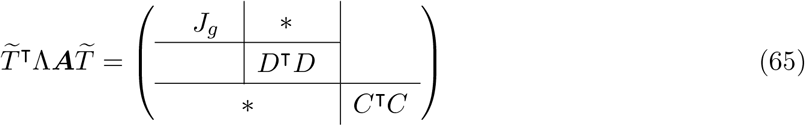

as desired. In the special case that dim coker ν = 0 (i.e., no conserved quantities), it is straightforward to see

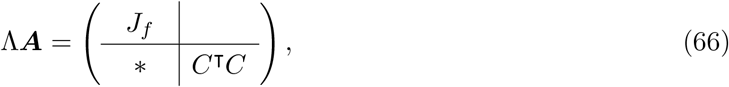

and then it is obvious that det Λ · det ***A*** = det (*C*^⊺^*C*) · det *J*_*f*_. ◼

### Corollary 1

*For any given state* ***x*** ∈ ℝ^*M*^, *we have*

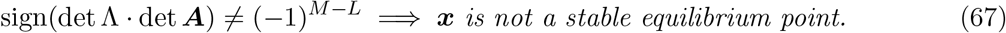

(proof) If ***x*** is a stable equilibrium point, then *J*_*g*_ is a real matrix such that each of (*M* − *L*) eigenvalues possesses a negative real part. With the complex conjugate root theorem and **Theorem 1**, one obtains

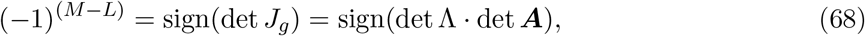

since the determinants of (*C*^⊺^*C*) and (*D*^⊺^*D*) must be positive. Hence, if the equation does not hold, then ***x*** is not a stable equilibrium point. ◼

## Appendix D: Code for numerical demonstrations

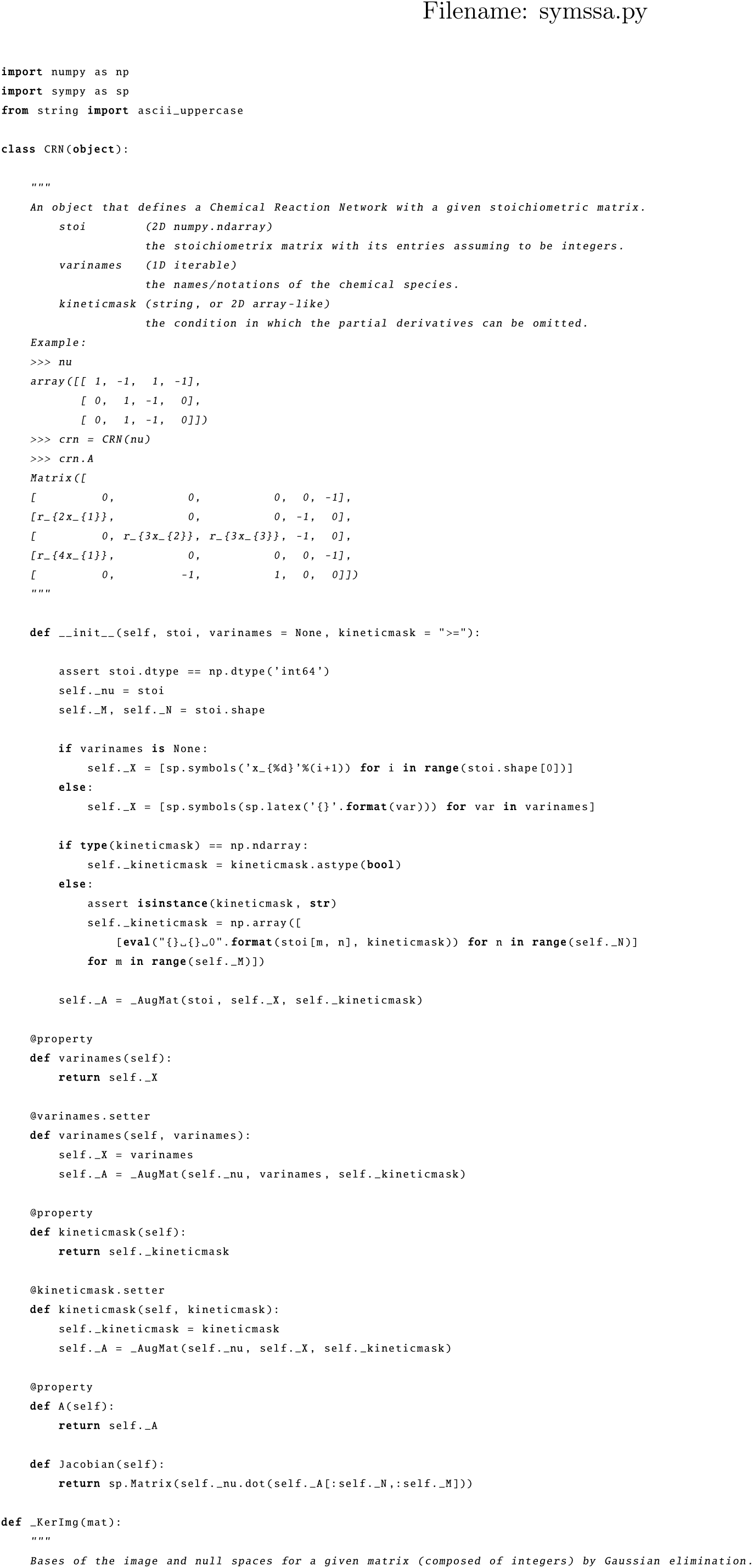
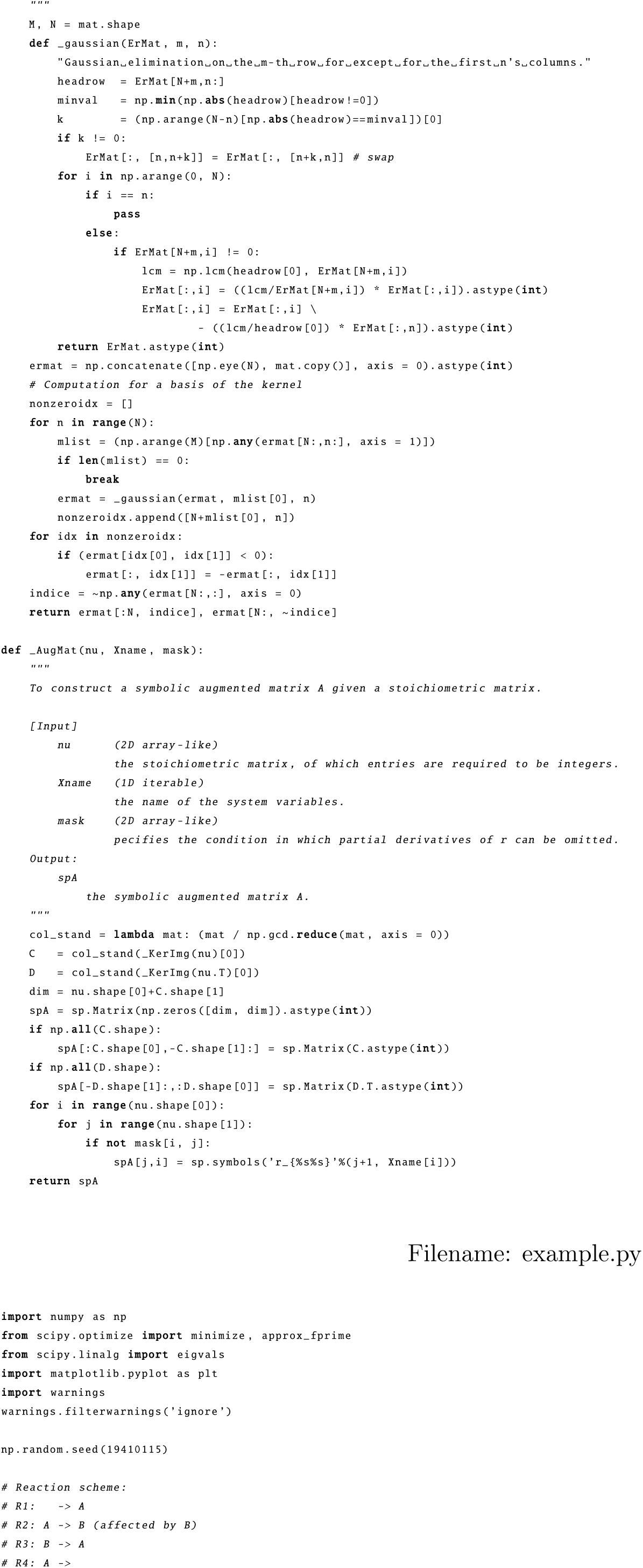
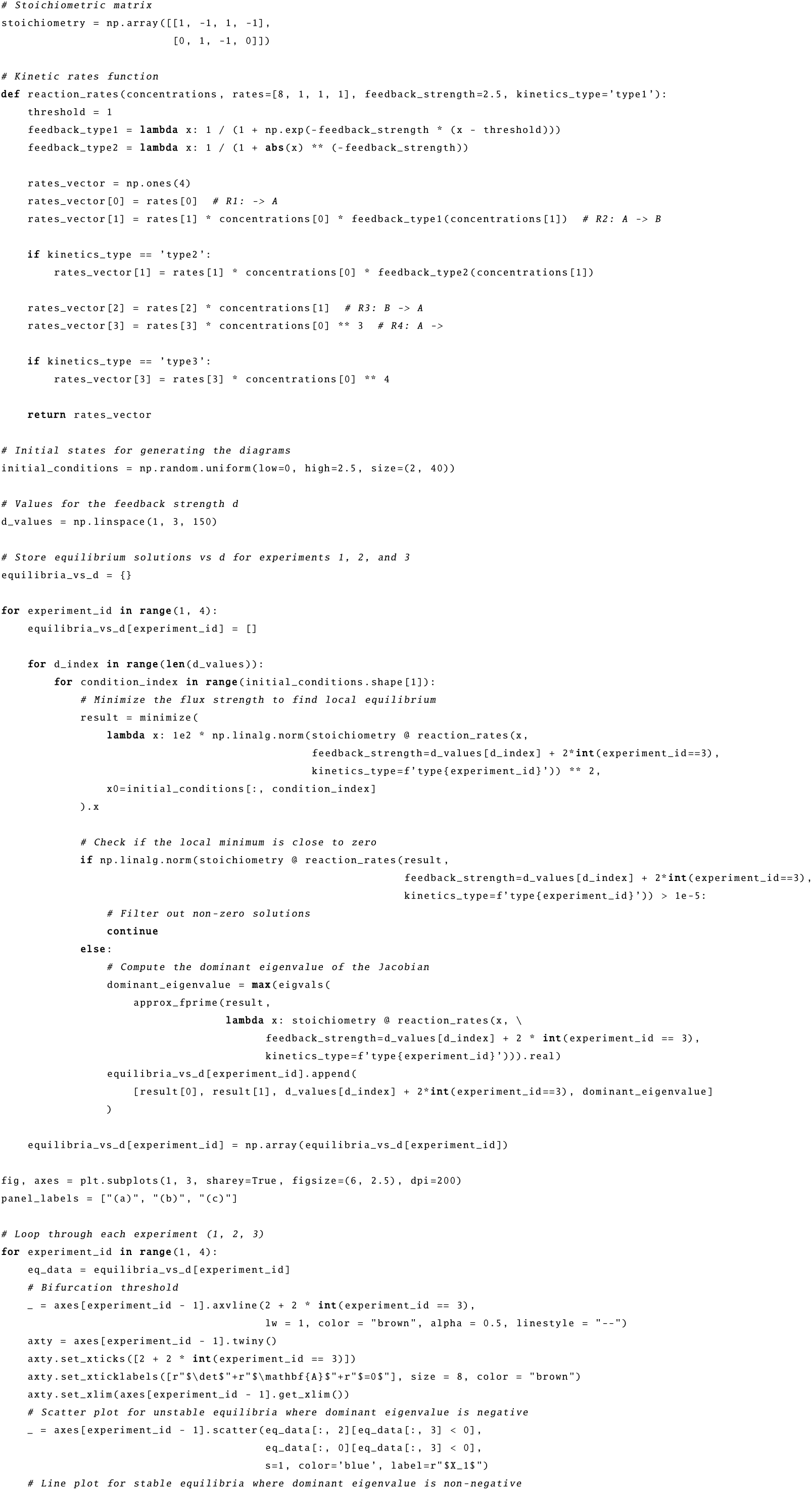
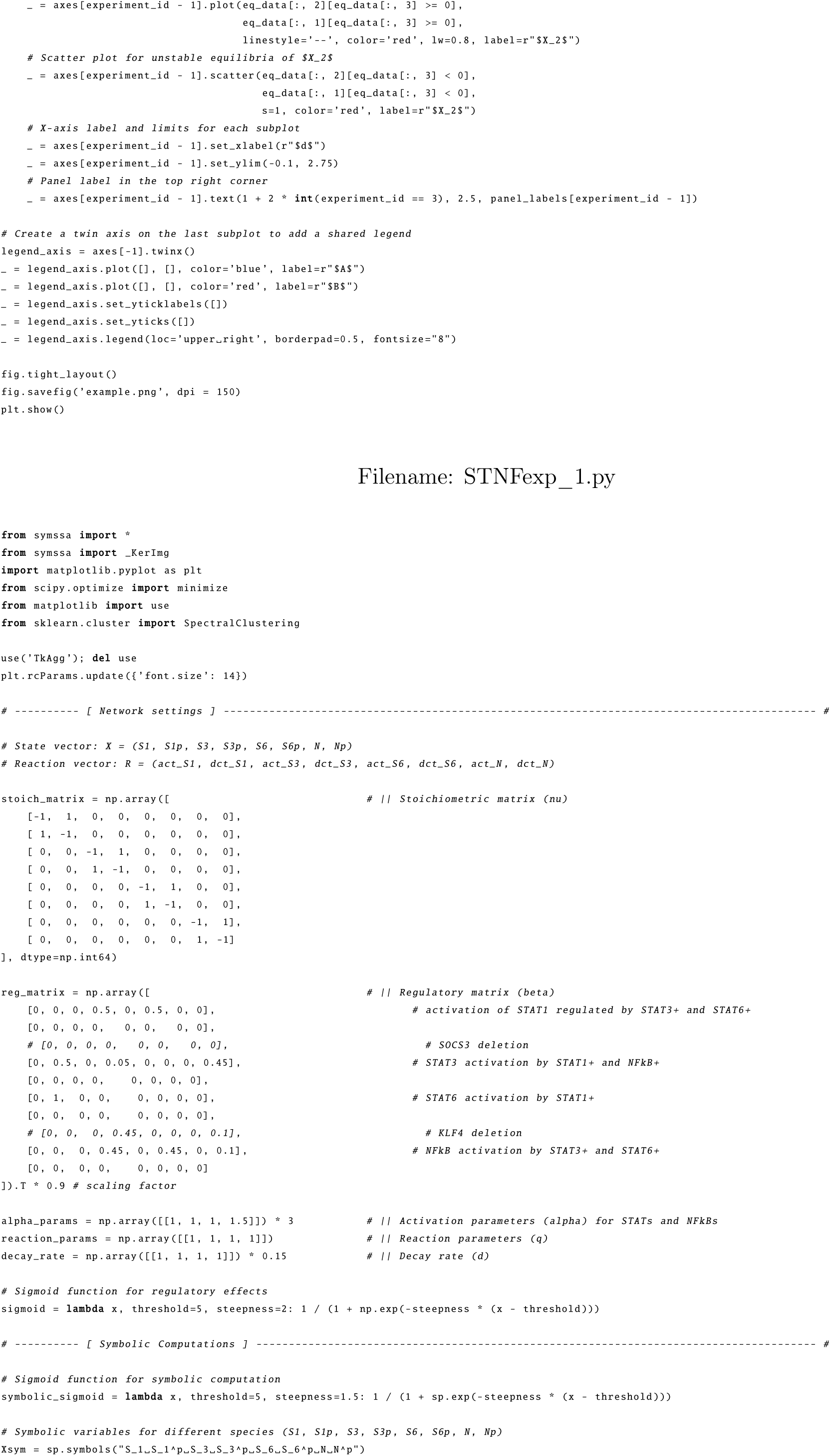
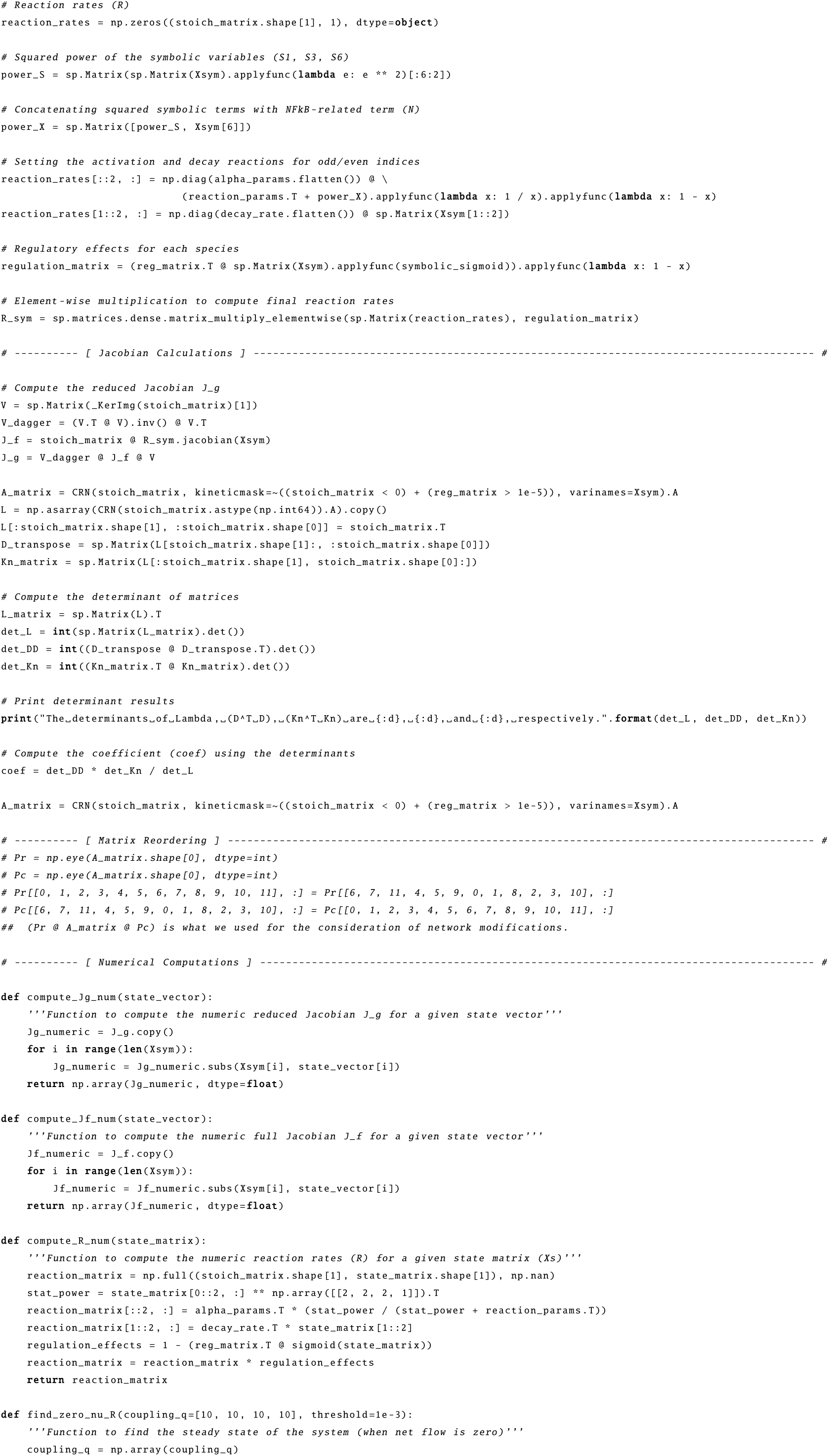
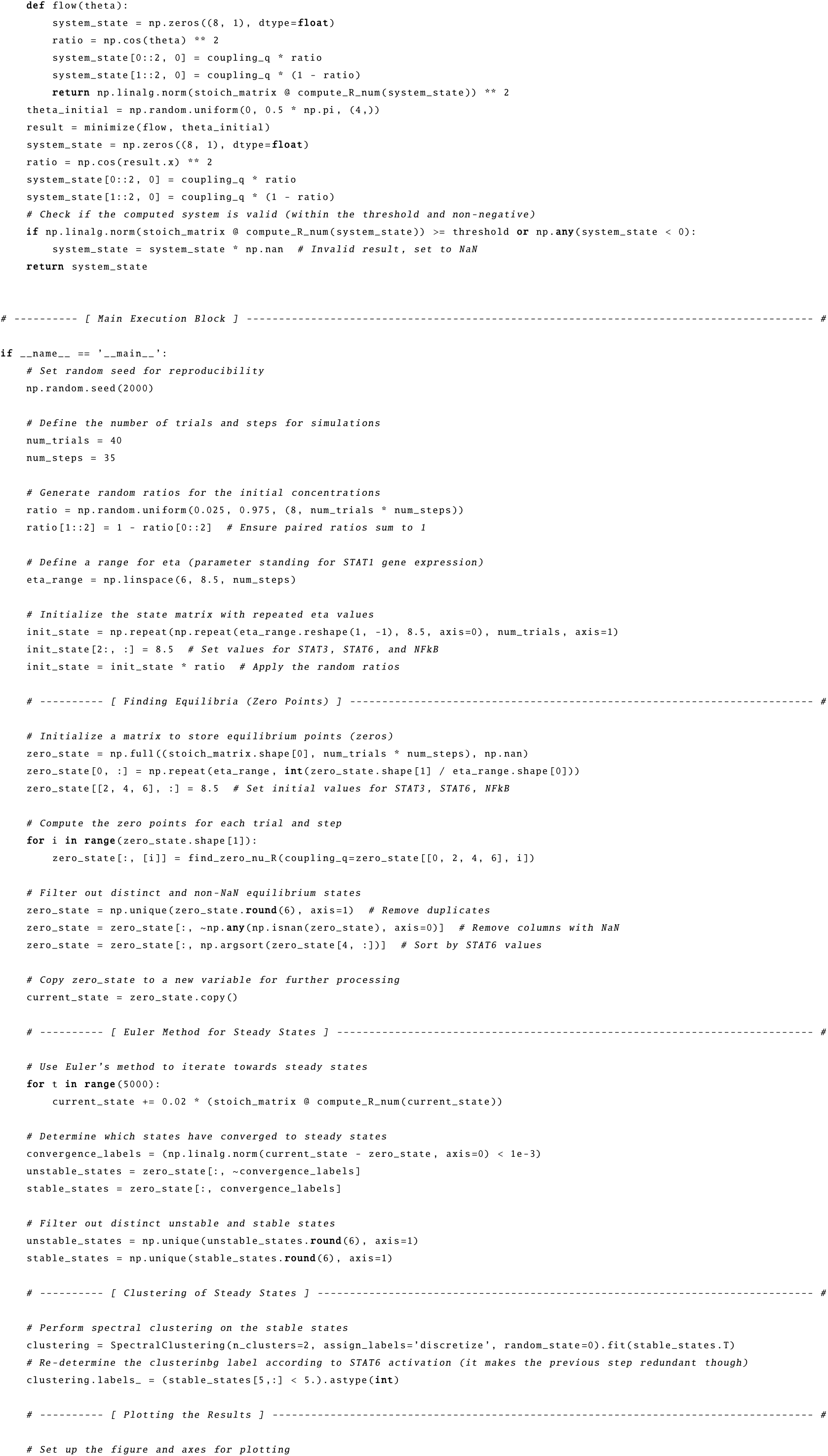
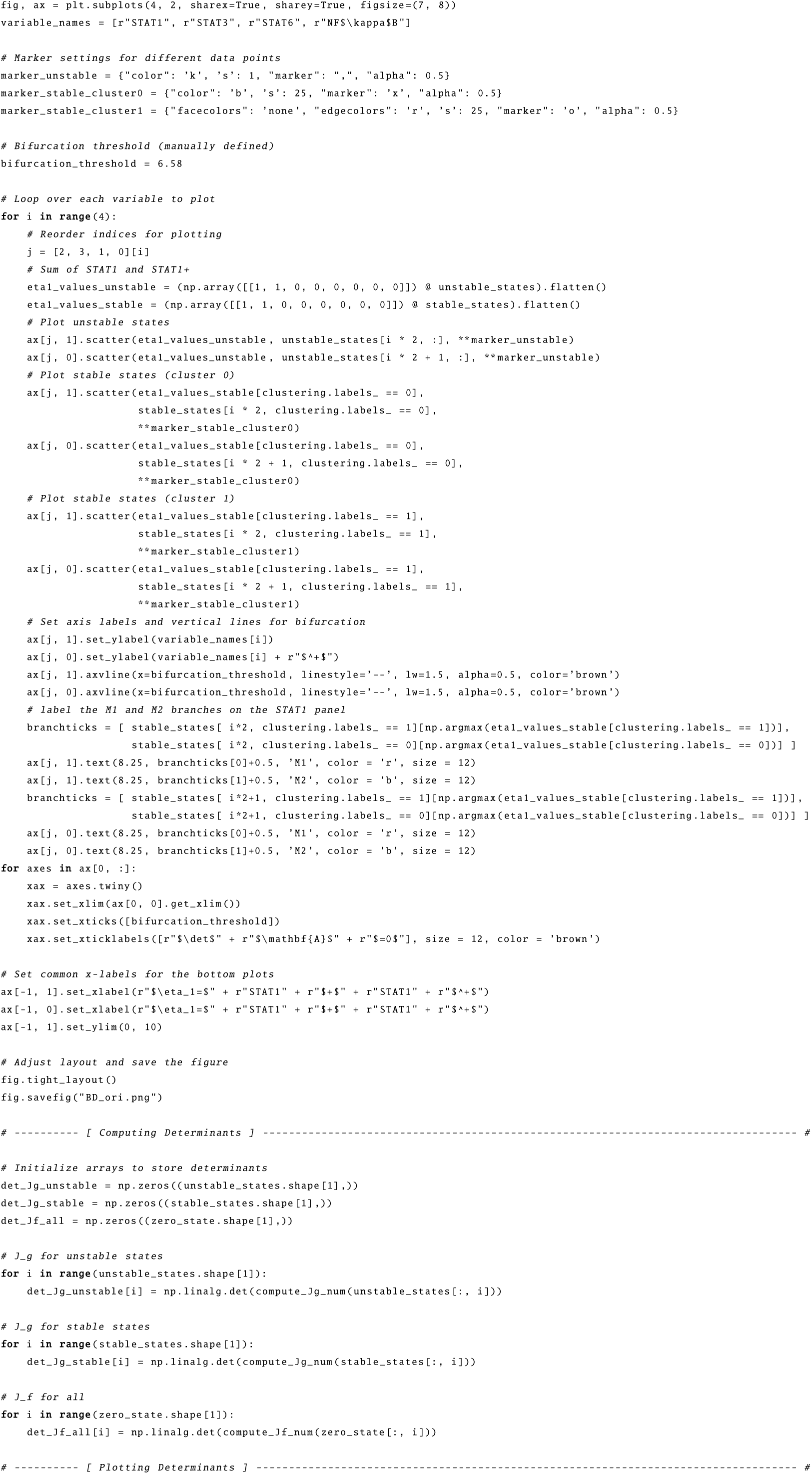
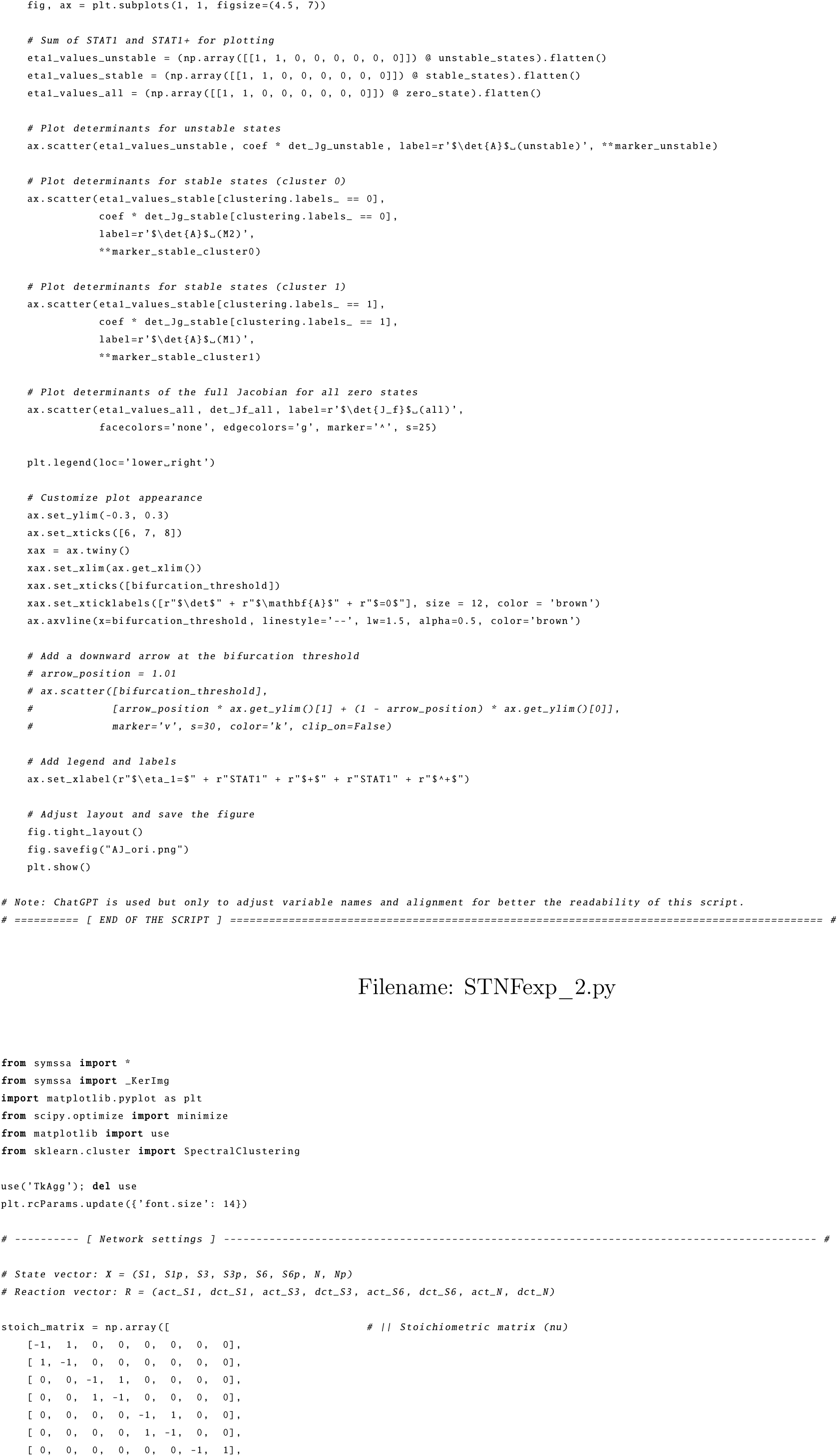
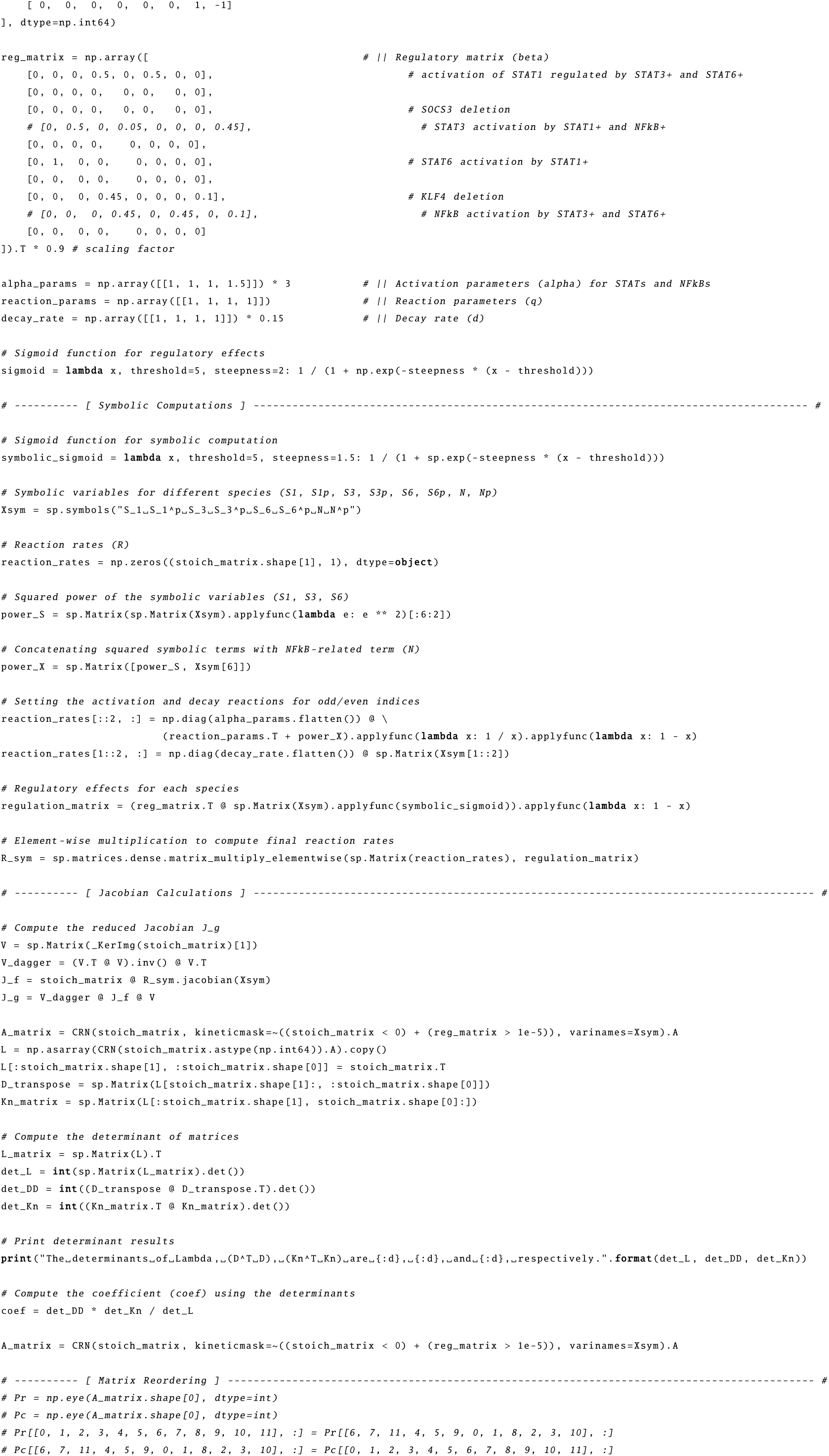
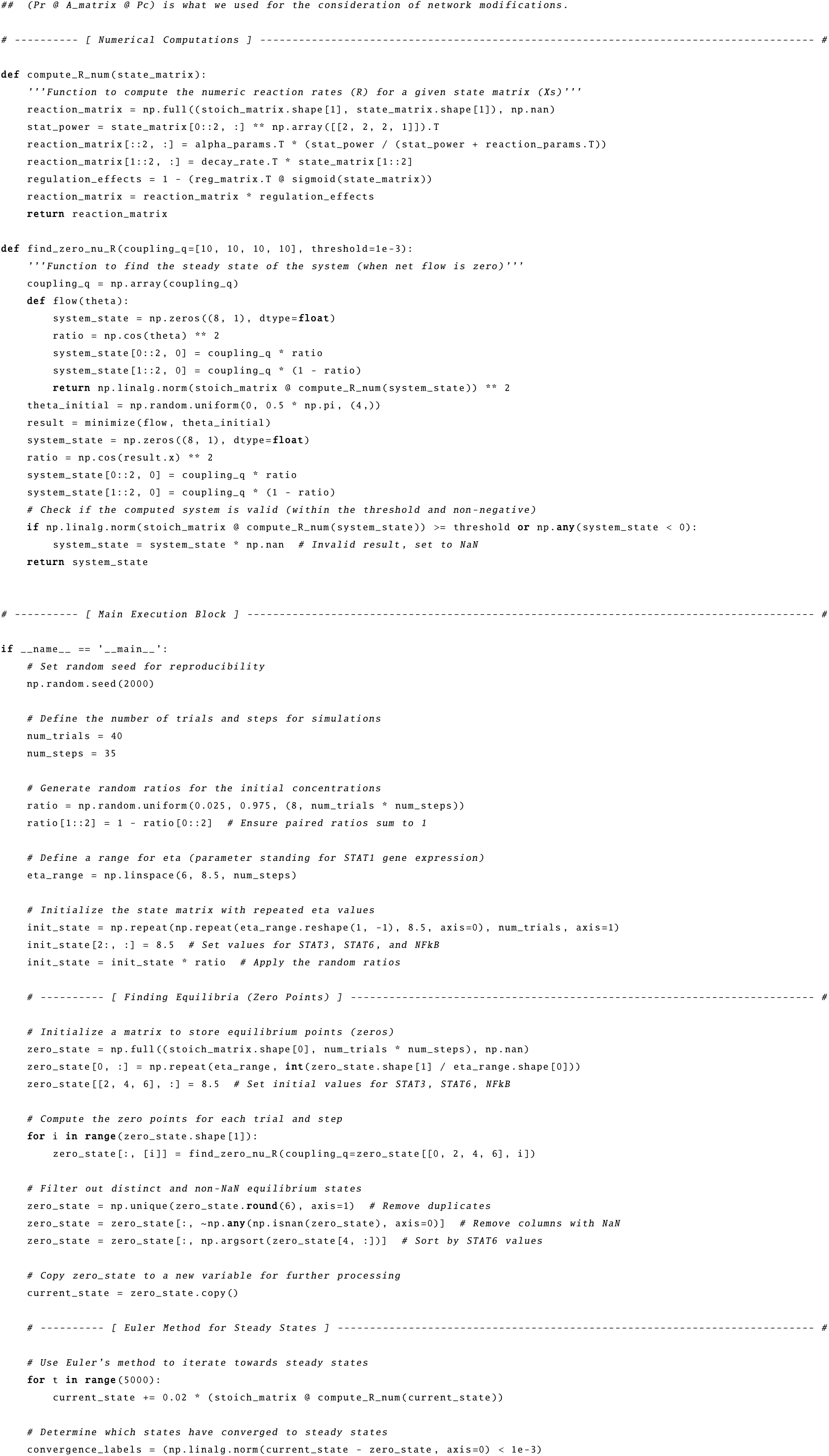
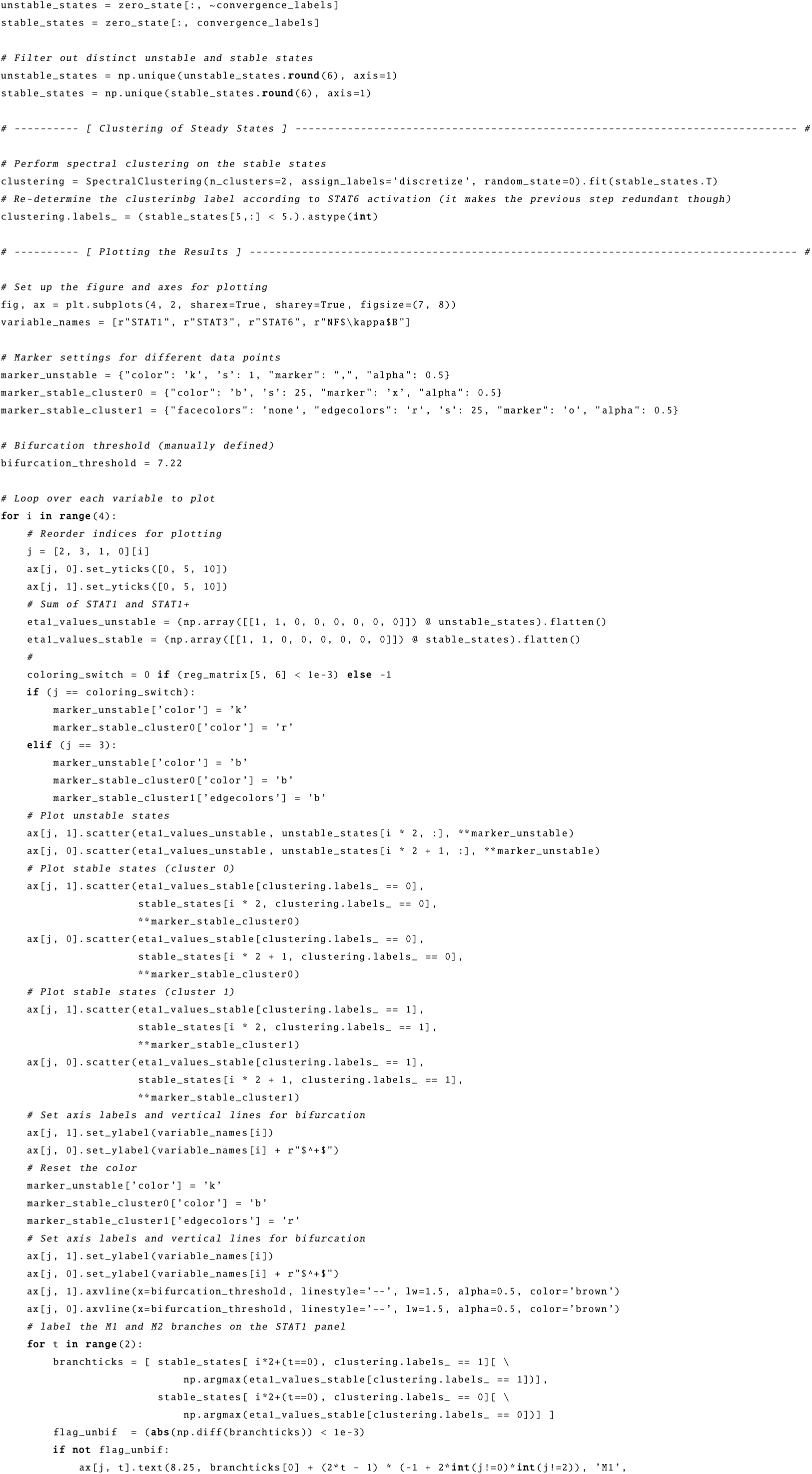
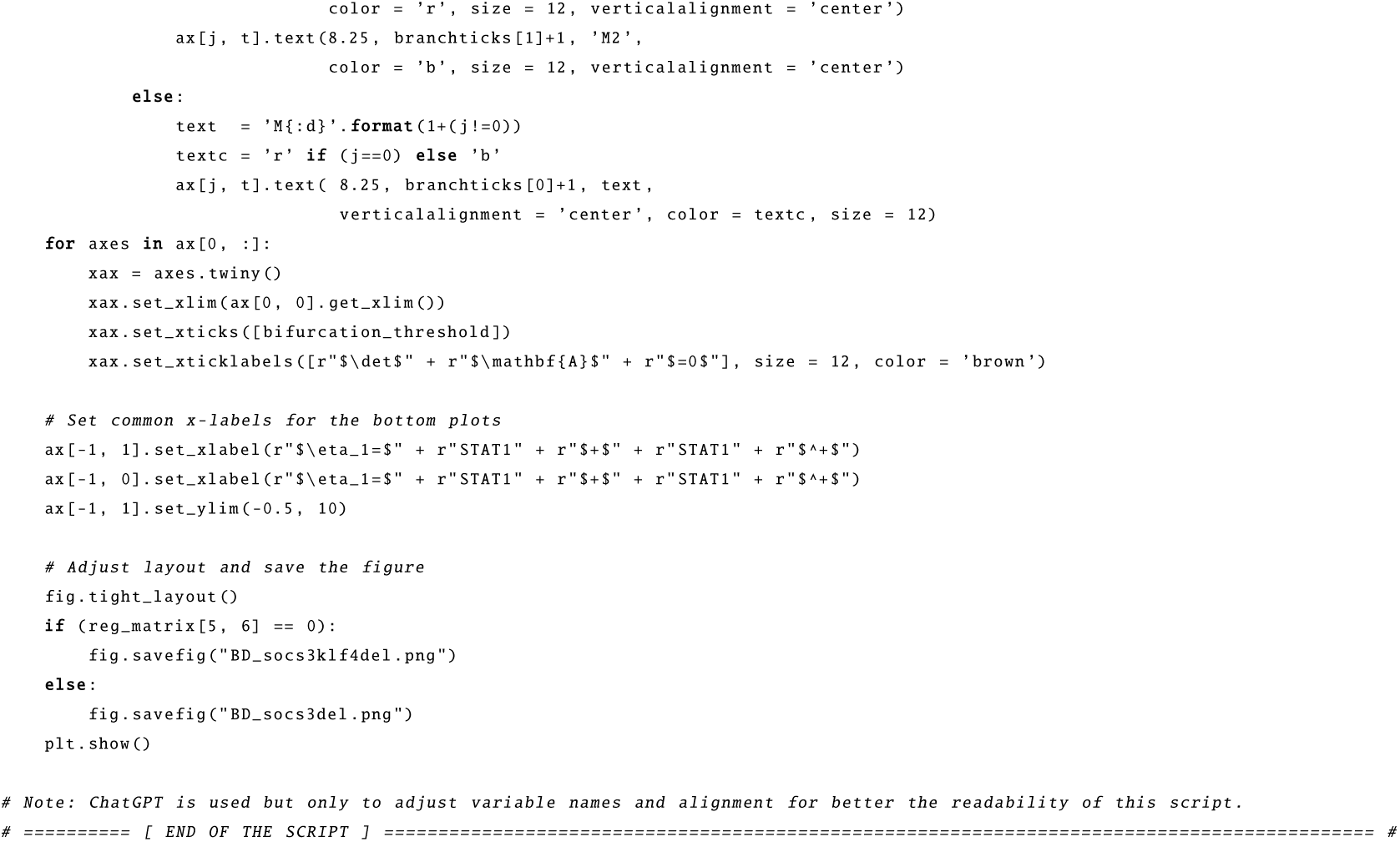

